# A novel *qacA* allele results in an elevated chlorhexidine glu conate minimum inhibitory concentration in cutaneous *Staphylococcus epidermidis* isolates

**DOI:** 10.1101/490805

**Authors:** Amin Addetia, Alexander L Greninger, Amanda Adler, Shuhua Yuan, Negar Makhsous, Xuan Qin, Danielle M Zerr

## Abstract

Chlorhexidine gluconate (CHG) is a topical antiseptic widely used in healthcare settings. In *Staphylococcus* spp., the pump QacA effluxes CHG, while the closely related QacB cannot due to a single amino acid substitution. We characterized 1,050 cutaneous *Staphylococcus* isolates obtained from 173 pediatric oncology patients enrolled in a multicenter CHG bathing trial. CHG susceptibility testing revealed 63 (6%) of these isolates had elevated CHG MICs (≥ 4 μg/mL). Screening of all 1,050 isolates for *qacA/B* by restriction fragment length polymorphism (RFLP) yielded 56 isolates with a novel *qacA/B* RFLP pattern, *qacAB*_*273*_. The CHG MIC was significantly higher for *qacAB*_*273*_-positive isolates (MIC_50_: 4 μg/mL, [range: 0.5 – 4 μg/mL]) compared to other *qac* groups: *qacA*-positive (n=559, 1 μg/mL, [0.5 – 4 μg/mL]), *qacB*-positive (n=17, 1 μg/mL, [0.25 – 2 μg/mL]), and *qacA/B-*negative (n=418, 1 μg/mL, [0.125 – 2 μg/mL], p=0.001). The *qacAB*_*273*_-positive isolates also displayed a high proportion of methicillin resistance (96.4%) compared to other *qac* groups (24.9 – 61.7%, p=0.001). Whole genome sequencing revealed that *qacAB*_*273*_-positive isolates encoded a variant of QacA with 2 amino acid substitutions. This new allele, named *qacA4*, was carried on the novel plasmid pAQZ1. The *qacA4*-carrying isolates belonged to the highly resistant *S. epidermidis* clone ST2 and were collected from multiple centers across the United States and Canada. Curing an isolate of *qacA4* resulted in a four-fold decrease in the CHG MIC, confirming the role of *qacA4* in the elevated CHG MIC. Our results highlight the importance of further studying *qacA4* and its functional role in clinical staphylococci.

**Importance:** *Staphylococcus epidermidis* is an important cause of infections in patients with implanted devices. Bathing with chlorhexidine gluconate (CHG), a topical antiseptic, has been shown to reduce rates of device-associated infections, especially those caused by *S. epidermidis*. In *S. epidermidis*, reduced susceptibility to CHG is associated with carriage of the *qacA* gene. As part of a multicenter CHG bathing trial, we obtained cutaneous *Staphylococcus* isolates from pediatric oncology patients across the United States and Canada. We identified a group of isolates capable of surviving in higher concentrations of CHG and determined a novel allele of *qacA*, termed *qacA4* and carried on the novel plasmid pAQZ1, was responsible for the isolates’ survival in higher CHG concentrations. The *qacA4*-carrying *S. epidermidis* isolates belonged to the highly resistant and virulent ST2 clonal type. Our results highlight the need to understand the global distribution of novel *qacA* alleles, including *qacA4*, and their mechanistic effect on efflux.

## Introduction

*Staphylococcus epidermidis* is a typical resident of the skin flora and an important cause of device-associated infections, especially central line associated bloodstream infections (1). The success of *S. epidermidis* as an opportunistic pathogen derives from its ability to bind indwelling devices through the formation of a biofilm (2–4) and the high rate of antimicrobial resistance within the population (5, 6).

With a favorable safety profile and broad-spectrum and residual activity (7), chlorhexidine gluconate (CHG), is a promising option for skin cleansing and antisepsis for the prevention of device-associated infections. Bathing with CHG has been demonstrated to reduce the rates of central-line associated bloodstream infections (8, 9), acquisition of multidrug resistant organisms (10), and blood culture contamination, which is frequently caused by *S. epidermidis* (11, 12). Furthermore, topical applications of CHG have been demonstrated to significantly reduce cutaneous microbial burden (13, 14). However, increasing usage of CHG may select for organisms with decreased susceptibility to CHG and increased resistance to commonly prescribed antimicrobials (13, 15–17).

In *Staphylococcus* spp., *qacA* encodes a 514-amino acid, 14 transmembrane segment pump with the capacity to efflux CHG (18–20). The pump encoded by the closely related *qacB* differs from *qacA* by only 7-9 nucleotides, but does not have the ability to efflux CHG (18, 21). A single nucleotide variant (SNV) (968C>A) resulting in a substitution, Ala323Asp, in transmembrane segment 10 accounts for the differing substrate specificities of QacA and QacB (18). Currently, three alleles of *qacA* have been described; however, no functional differences between the pumps encoded by these three alleles have been reported (21).

Beyond CHG, QacA is responsible for the efflux of a broad range of mono- and divalent cations, including dyes and quaternary ammonium compounds (20). In *S. epidermidis*, *qacA* is most frequently carried by the plasmid pSK105, which also carries the aminoglycoside resistance gene *aacA-aphD* (22). Other plasmids carrying *qacA* may contain the trimethoprim resistance gene *dfrA*, the *blaZ β*-lactamase, or genes encoding heavy metal efflux pumps (22).

In addition to QacA, the 107-amino acid, 4 transmembrane segment efflux pump encoded by *smr*, also known as *qacC*, has been implicated in the efflux of CHG (23–25). While unrelated to QacA and QacB, Smr demonstrates the capacity to efflux a similar, yet narrower range of monovalent cations (24, 25).

In our study, cutaneous *Staphylococcus* isolates were obtained from pediatric oncology patients enrolled in a multicenter randomized controlled CHG bathing trial. We identified a subpopulation of isolates with an elevated CHG MIC, which we defined as an MIC ≥ 4 μg/mL. To investigate the genetic basis of the elevated CHG MIC, we screened the isolates for the *qacA/B* genes via PCR and restriction fragment length polymorphism (RFLP). From this screening, we identified a previously undescribed RFLP pattern, termed *qacAB*_*273*_, in a subset of isolates. We then determined whether the *qacAB*_*273*_ RFLP pattern was associated with a significantly higher CHG MIC when compared to the *qacA*-positive, *qacB*-positive, and *qacA/B*-negative isolates. We also described the sequence of the novel *qacA* allele, referred to as *qacA4*, producing the novel *qacAB*_*273*_ RFLP pattern and characterized the isolates carrying *qacA4*. Furthermore, through curing experiments, we investigated the role of *qacA4* in causing elevated CHG MICs in *S. epidermidis*.

## Results

### Overview of study population

In total, 1050 cutaneous *Staphylococcus* isolates were obtained from 173 patients. The study isolates primarily consisted of coagulase negative *Staphylococcus* with *S. epidermidis* being the most frequently recovered species (53.1%), while *S. aureus* accounted for just 2.9% of the study population (Table 1). In addition to *S. epidermidis*, 17 other coagulase negative *Staphylococcus* species were identified in the study population. Of note, four coagulase negative *Staphylococcus* isolates could not be speciated by MALDI-TOF.

**Table 1.**
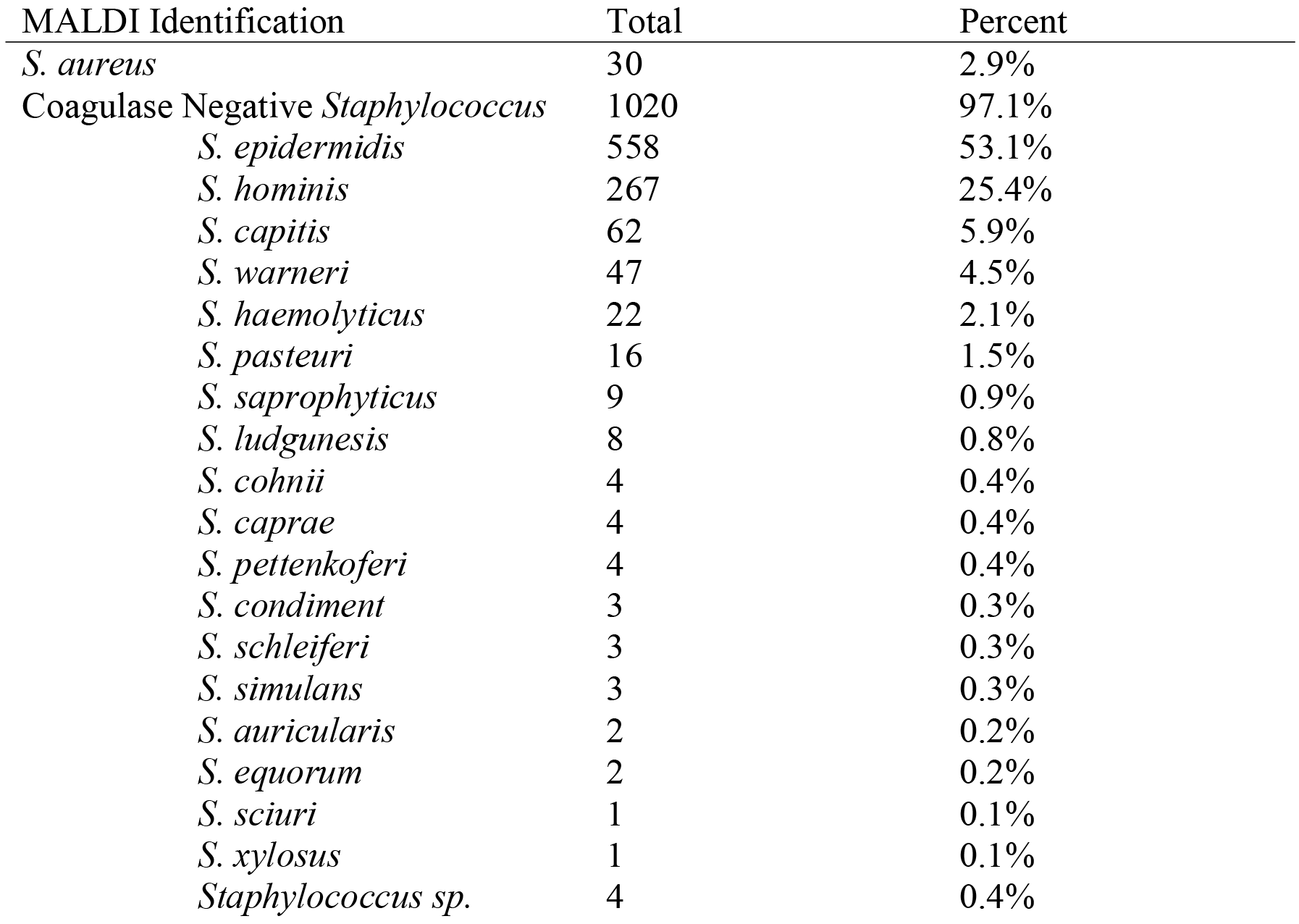
Overview of the cutaneous *Staphylococcus* isolates (n = 1050) included in this study.

### A subset of Staphylococcus isolates have an elevated CHG MIC

Measuring the CHG MICs across all 1050 isolates yielded 63 isolates with elevated CHG MICs, defined as an MIC ≥ 4 μg/mL (Figure 1a). All of these isolates were identified as *S. epidermidis*.

**Figure 1.**
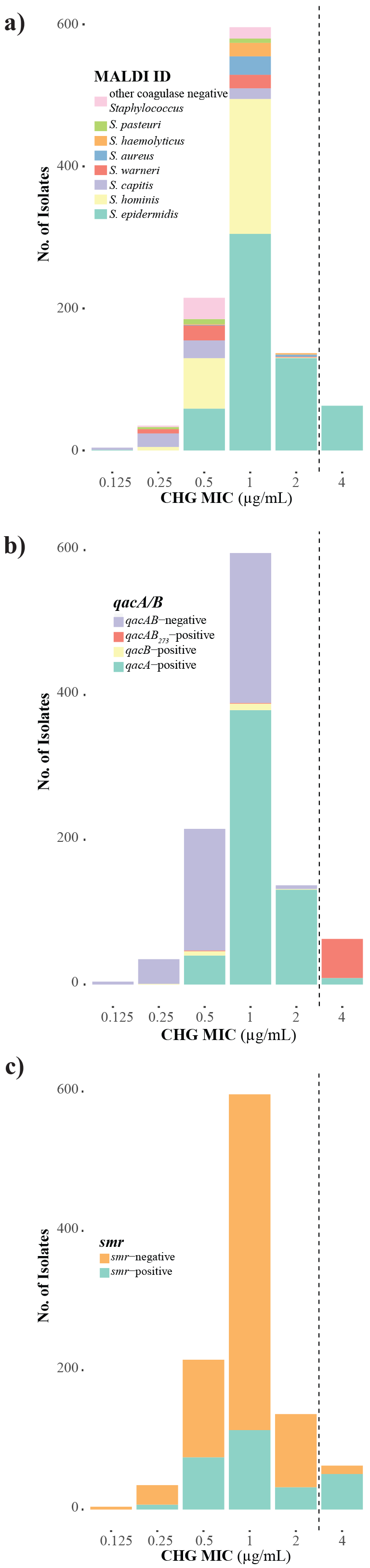
CHG MIC distribution of the 1050 cutaneous *Staphylococcus* isolates included in our study grouped by a) species, b) *qacA/B* PCR and RFLP patterns, and c) *smr* PCR results. The dashed line indicates the concentration we defined as an elevated CHG MIC (≥ 4 μg/mL).

### Identification of a novel qacA/B RFLP pattern

Isolates were screened for the *qacA/B* genes to explore the genetic basis of the elevated CHG MICs. PCR amplification of the *qacA/B* gene resulted in an 864 bp product. Digestion of the *qacA/B* PCR product with AluI resulted in the presence of a characteristic 198 bp fragment for *qacA*-positive isolates and a characteristic 165 bp fragment for *qacB*-positive isolates. A third subpopulation of isolates was distinguished by the appearance of a 273 bp fragment (Figure 2), and is referred to as *qacAB*_*273*_-positive isolates hereafter.

**Figure 2.**
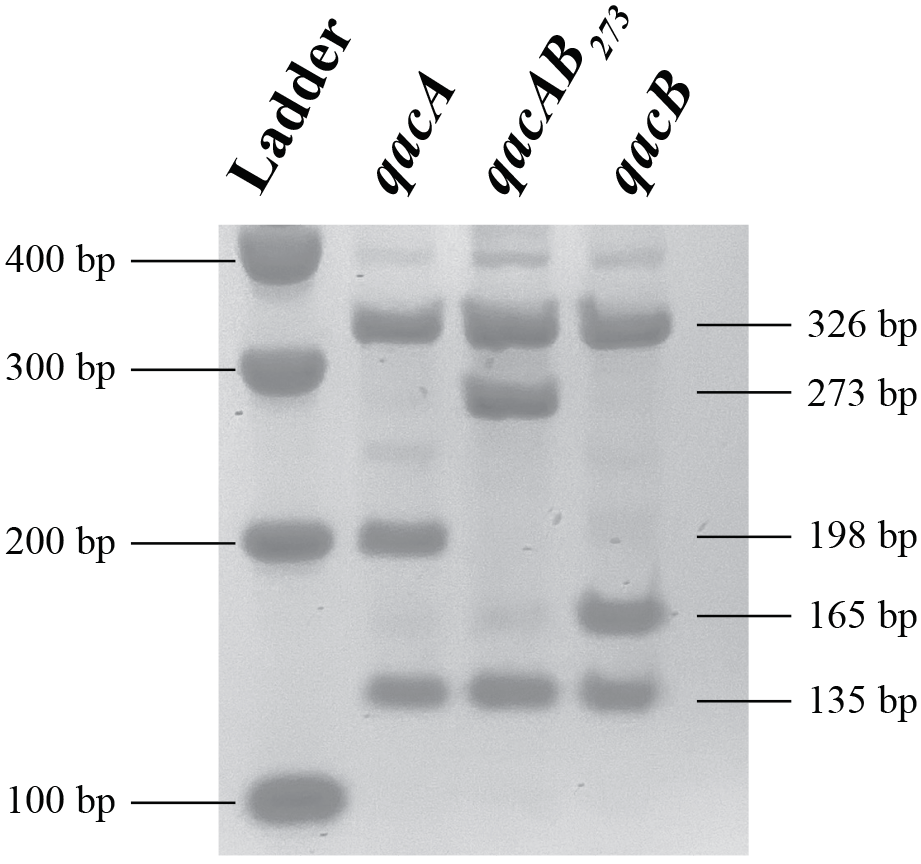
RFLP patterns observed from the AluI restriction digest of the *qacA/B* PCR amplicon. Isolates were classified as *qacA*-postive, *qacAB*_*273*_-positve, or *qacB*-positive based on the presence of a 198 bp, 273 bp, or 165 bp fragment, respectively (Alam et al., 2003). Ladder: 100 bp (Promega).

Of the 1050 isolates, 632 contained a *qacA/B* gene as identified by PCR. Based on the results of the RFLP analysis, 559 were classified as *qacA*-positive, 17 as *qacB*-positive, and 56 as *qacAB*_*273*_-positive (Table 2). The *qacA/B* genes were detected in 8 different coagulase-negative *Staphylococcus* species. When screened for carriage of *smr*, 279 of the 1050 isolates were classified as *smr*-positive (Table 2). In total, 12 unique coagulase-negative *Staphylococcus* species carried *smr*. Notably, the *qacA/B* genes and *smr* were not detected in any of the *S. aureus* isolates.

**Table 2.** Comparison of the CHG MIC distribrutions, measured in μg/mL, of the *qacA*-positive, *qacB*-positive, *qacAB*_*273*_-positive, and *qacA/B*-negative isolates and the CHG MIC distributions of the *smr*-positive and *smr*-negative isolates.

|  | *qacA* | *qacB* | *qacAB*<sub>273</sub> | *qacA/B* – | *smr* + | *smr* – |
| --- | --- | --- | --- | --- | --- | --- |
| Total Isolates | 559 | 17 | 56 | 418 | 279 | 771 |
| MIC<sub>50</sub> | 1 | 1 | 4 | 1 | 1 | 1 |
| MIC<sub>90</sub> | 2 | 1 | 4 | 1 | 2 | 1 |
| Range | 0.5 – 4 | 0.25 – 2 | 0.5 – 4 | 0.125 – 2 | 0.25 – 4 | 0.125 – 4 |

### The qacAB_273_ RFLP pattern is associated with an elevated CHG MIC

Next, the relationship between elevated CHG MICs and detection of the *qacA/B* and *smr* genes was examined. A *qacA/B* gene was detected in each of the 63 isolates with an elevated CHG MIC: 54 were classified as *qacAB*_*273*_-positive and 9 as *qacA*-positive (Figure 1b). None of the isolates with an elevated CHG were classified as *qacB*-positive. Furthermore, 51 of the 63 isolates with elevated CHG were categorized as *smr*-positive and the remaining 12 as *smr*-negative (Figure 1c).

To further investigate if the *qacAB*_*273*_ RFLP pattern was associated with an elevated CHG MIC, differences in the CHG MIC distributions of the *qacA/B* containing isolates were assessed. The CHG MIC was significantly higher for the *qacAB*_*273*_-positive isolates as compared to the *qacA*-positive, *qacB*-positive, and *qacA/B*-negative isolates (p = 0.001); the results did not change when restricting the analyses to one randomly chosen isolate per patient per *qacA/B* group (Table 2). In addition, the CHG MIC distributions of the *smr*-positive and *smr*-negative isolates were compared. The CHG MIC was significantly higher for the *smr*-positive isolates compared to the *smr*-negative isolates (p=0.02); however, this comparison was no longer significant when the analyses were restricted to one randomly chosen isolate per patient per *smr* group as one individual accounted for 20% of the *smr*-positive isolates with elevated MICs (p=0.11) (Table 2).

Additionally, CHG MIC distributions associated with *qacA/B* and *smr* resistance gene combinations among all isolates were assessed to determine if a particular resistance gene combination was associated with elevated CHG MICs. This comparison revealed *qacAB*_*273*_ rather than a particular resistance gene combination was associated with elevated CHG MICs (p=0.001); the results did not change when restricting the analyses to one randomly chosen isolate per patient per gene combination (Table 3).

**Table 3.** Comparison of the CHG MIC distributions, measured in μg/mL, associated with the eight CHG resistance gene combinations detected in our isolates.

|  | *qacA*<br>*smr* + | *qacA*<br>*smr* – | *qacB*<br>*smr* + | *qacB*<br>*smr* – | *qacAB*<sub>273</sub><br>*smr* + | *qacAB*<sub>273</sub><br>*smr* – | *qacA/B* –<br>*smr* + | *qacA/B* –<br>*smr* – |
| --- | --- | --- | --- | --- | --- | --- | --- | --- |
| Total Isolates | 101 | 458 | 5 | 12 | 50 | 6 | 123 | 295 |
| MIC<sub>50</sub> | 1 | 1 | 1 | 0.75 | 4 | 4 | 0.5 | 1 |
| MIC<sub>90</sub> | 2 | 2 | 1.6 | 1 | 4 | 4 | 1 | 1 |
| Range | 0.5 – 4 | 0.5 – 4 | 0.5 – 2 | 0.25 – 1 | 4 – 4 | 0.5 – 4 | 0.25 – 1 | 0.125 – 2 |

The *qacAB*_*273*_-positive isolates exhibited higher rates of resistance to methicillin (96.4%) and other commonly prescribed antimicrobials, including erythromycin (ERY, 92.9%), ciprofloxacin (CIP, 96.4%), gentamicin (GEN, 89.3%), and sulfamethoxazole/trimethoprim (SXT, 98.2%), as compared to the *qacA*-positive, *qacB*-positive and *qacA/B*-negative isolates (p<0.001 for all comparisons); the results did not change when restricting analyses to one randomly chosen isolate per patient per *qacA/B* group (Table 4). All *qacA/B* groups exhibited rates of resistance less than 1% to linezolid (LZD), rifampin (RIF), and vancomycin (VAN).

**Table 4.** Comparison of the proportion of *qacA*-positive, *qacB*-positive, *qacAB*_*273*_-positive, and *qacA/B*-negative isolates resistant to commonly prescribed antimicrobials: cefoxitin (FOX), erythromycin (ERY), ciprofloxacin (CIP), gentamicin (GEN), sulfamethoxazole/trimethoprim (SXT), linezolid (LZD), rifampin (RIF), and vancomycin (VAN).

| Antimicrobial | *qacA* | *qacB* | *qacAB*<sub>273</sub> | *qacA/B* – |
| --- | --- | --- | --- | --- |
| FOX* | 61.7 | 52.9 | 96.4 | 22.7 |
| ERY* | 76.0 | 88.2 | 92.9 | 30.1 |
| CIP* | 25.0 | 5.9 | 96.4 | 8.4 |
| GEN* | 15.0 | 0 | 89.3 | 2.6 |
| SXT* | 60.1 | 35.3 | 98.2 | 22.7 |
| LZD | 0 | 0 | 0 | 0 |
| RIF | 0.9 | 0 | 0 | 0 |
| VAN | 0 | 0 | 0 | 0 |

\* P-value <0.001 for all comparisons. Results were generated using all isolates; results did not change when restricting analyses to one randomly chosen isolate per patient per *qacA/B* group.

### Whole genome sequencing of qacA/B-positive isolates yields novel qacA alleles

To further investigate the *qacA/B* gene in the *qacAB*_*273*_-positive isolates, the genomes of 9 *qacAB*_*273*_-positive *S. epidermidis* isolates were compared to the genomes of 10 *qacA*-positive and 4 *qacB*-positive *S. epidermidis* isolates (Table S1). All 9 of the *qacAB*_*273*_-positive isolates had elevated CHG MICs, while none of the 10 *qacA*-positive and 4 *qacB*-positive isolates had elevated CHG MICs.

The sequence of *qacA/B* gene was highly conserved in the 9 *qacAB*_*273*_-positive isolates with elevated CHG MICs. As the *qacA/B* gene of the *qacAB*_*273*_-positive isolates contained the distinguishing *qacA* nucleotide 968A, the gene was classified as a novel allele of *qacA*. As shown in Figure 3a, this allele contained three SNVs (470C>G, 819G>A, and 1133C>T) compared to a reference *qacA* sequence (AB566410), and is referred to as *qacA4* (MK040360) henceforth. The SNV at position 1133 in *qacA4* resulted in the loss of an AluI digestion site, explaining the novel RFLP pattern observed in Figure 2. Two of the SNVs resulted in amino acid substitutions, Ala157Gly and Ala378Val, in transmembrane segments 5 and 12, respectively (Figure 3b).

**Figure 3.**
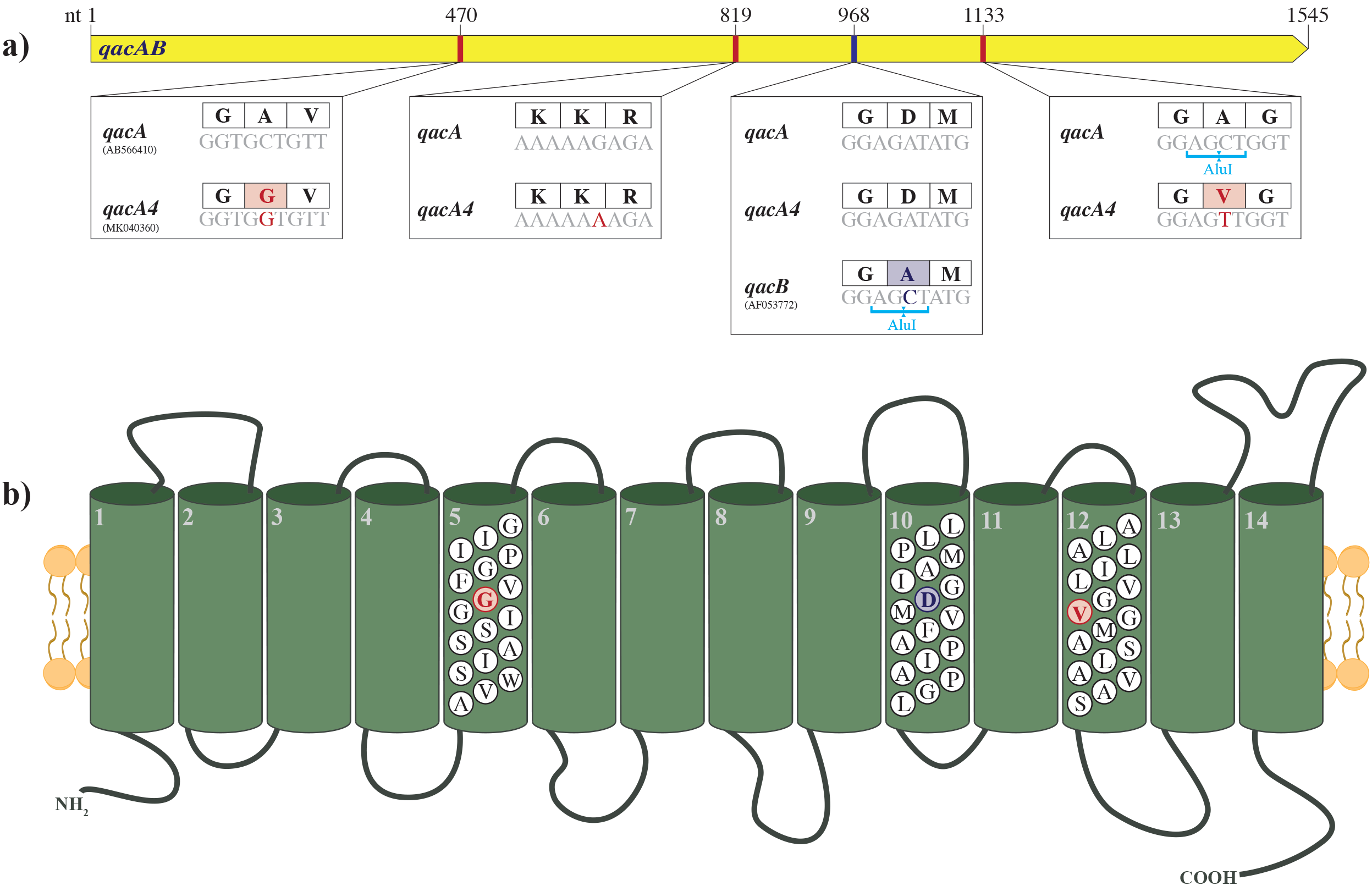
Comparison of a) the sequence of *qacA4* (MK040360), a reference *qacA* sequence (AB566410), and a reference *qacB* sequence (AF053772). The associated AluI restriction sites are shown below the nucleotide sequences and the corresponding amino acid sequences are displayed in the boxes above the nucleotide sequences. Structure of b) the predicted efflux pump encoded by *qacA4*, adapted from previous representations of QacA (Paulsen et al., 1996) (Brown et al., 1998). The residues which distinguish QacA4 from the reference QacA are highlighted in red. Those which distinguish QacA from QacB are displayed in blue.

When compared to the three previously characterized alleles of *qacA* (*qacA1* (GU565967), *qacA2* (21), and *qacA3* (MK040360)) and the *qacA* alleles of the 10 *qacA*-positive *S. epidermidis* isolates, *qacA4* differed from these sequences by at least three SNVs – including all three that distinguished *qacA4* from the reference *qacA* sequence (Figure S1). Notably, from the 10 *qacA*-positive isolates we sequenced, we identified 5 additional novel *qacA* alleles: *qacA7* (MK040363), *qacA8* (MK040364), *qacA9* (MK040365), *qacA*10 (MK040366), and *qacA11* (MK040367) (Table S2; Figure S1). Similarly, comparing the sequence of *qacA4* to the other sequences of *qacA* deposited in NCBI GenBank further confirmed the SNVs at the three positions described above were unique to *qacA4*.

We also sequenced the genomes of the two *qacAB*_*273*_-positive isolates that did not have elevated CHG MICs (MIC < 4 μg/mL) to investigate their discordant genotypic-phenotypic relationship (Table S1). Each of the *qacA/B* genes in these two *qacAB*_*273*_-positive isolates lacked the three identifying *qacA4* mutations and differed from the reference *qacA* sequence by six SNVs (Figure 4). These SNVs resulted in six and seven amino acid substitutions compared to the reference *qacA* and *qacA4*, respectively (Figure 4). The *qacA/B* genes in these isolates were classified as two additional alleles of *qacA*, referred to as *qacA5* (MK040361) and *qacA6* (MK040362). Due to a SNV at position 1132, which resulted in the loss of an AluI digestion site, *qacA5* and *qacA6* displayed identical digestion patterns to *qacA4*.

**Figure 4.**
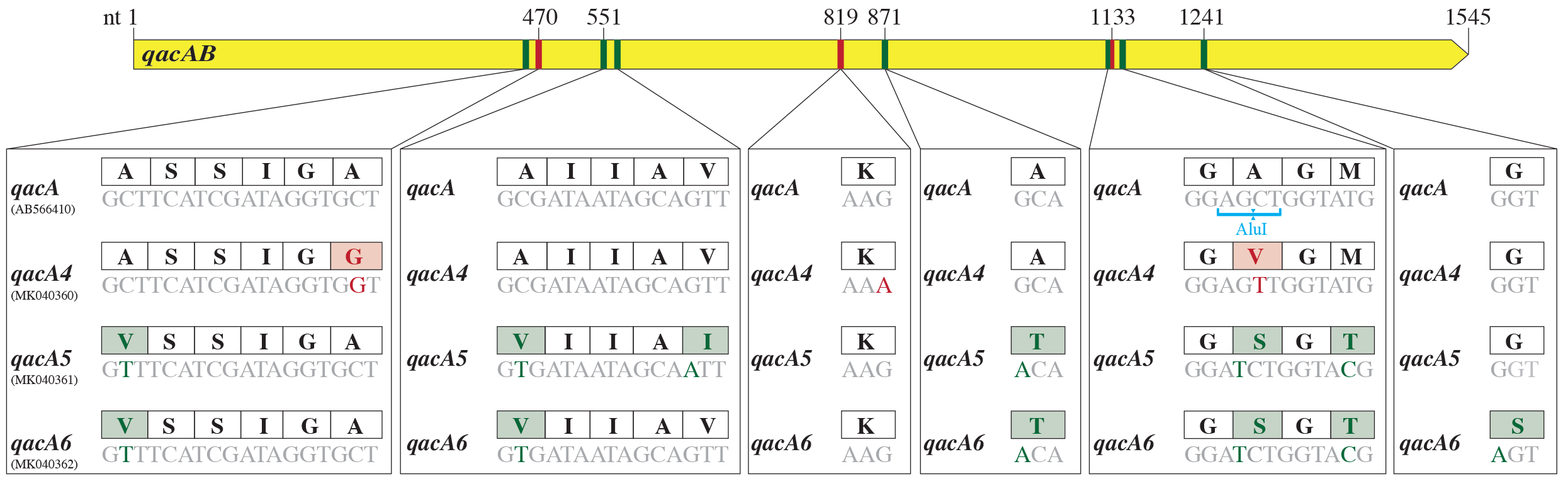
Comparison of the sequences of *qacA4* (MK040360), *qacA5* (MK040361), and *qacA6* (MK040362) and the reference *qacA* sequence (AB566410). The associated AluI restriction sites are shown below the nucleotide sequences and the corresponding amino acid sequences are displayed in the boxes above the nucleotide sequences. The nucleotides which distinguish *qacA4* from the reference *qacA*, *qacA5*, and *qacA6* are highlighted in red. Those which distinguish *qacA5* and/or *qacA6* from *qacA4* and the reference *qacA* are displayed in green.

We next sequenced genomes from the nine *qacA*-positive isolates that had elevated CHG MICs. From these isolates, we identified 4 *qacA* alleles: *qacA10, qacA12*, *qacA13*, and *qacA14* (Table S2; Figure S2). The sequences of the *qacA* genes in these isolates differed from the reference *qacA* sequence by 1 to 5 amino acid substitutions, and from *qacA4* by 2 to 8 amino acid substitutions (Figure S2). The allele *qacA14*, identified in isolate 96.5, contained one of the distinguishing coding changes of *qacA4*, Ala157Gly, but, not the other coding change. This allele encoded a unique amino acid substitution Pro328Leu, which distinguished the allele from the reference *qacA* and *qacA4*. Another isolate, 86.4, with an elevated CHG MIC carried the same *qacA10* allele as isolate 110.3, which did not have an elevated CHG MIC. The amino acid substitutions in these novel *qacA* alleles occurred in transmembrane segments 5, 6, 9, 10, 12, and 13 and in the extracellular loop between transmembrane segments 5 and 6.

### Identification of the novel resistance plasmid pAQZ1 containing qacA4 allele

The genomic context of the *qacA4* allele in *de novo* assemblies of two separate *qacAB*_*273*_-positive isolates with high coverage, isolates 91.2 and 107.2, was examined to understand whether *qacA4* was encoded chromosomally or on a plasmid. Both isolates carried *qacA4* on a 29,431 bp circular contig with coverage that was 2.8X higher than average chromosomal coverage, consistent with it being a plasmid (Figure 5a). The circular nature of the contig was verified by conducting PCR across the predicted junction site (data not shown). The plasmid, designated pAQZ1 (MK046687) henceforth, carrying *qacA4* contained the RepA replication initiation protein with a RepA_N domain (pfam06970). Similar to other RepA_N family plasmids (26, 27), the origin of replication of pAQZ1 is likely contained within *repA*. The plasmid also carried several genes involved in heavy metal efflux including, *copZ, copA*, and *czcD*, the *knt* kanamycin resistance gene, the *ble* bleomycin resistance gene, and an incomplete β-lactamase operon.

**Figure 5.**
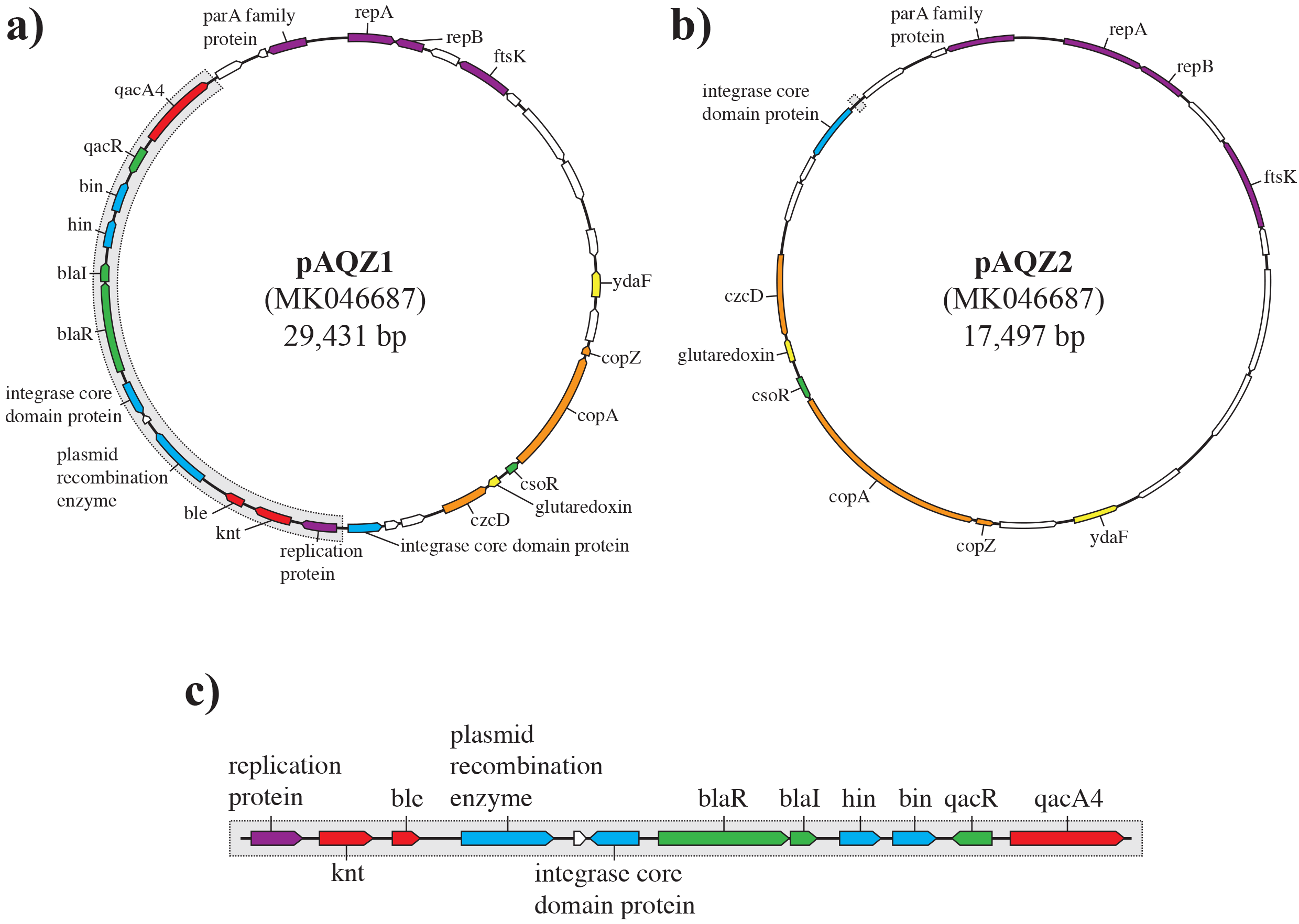
Schematic representations of a) the plasmid pAQZ1 containing *qacA4* obtained from the *de novo* assemblies of isolates 91.2 and 107.2, b) the plasmid pAQZ2 retained in isolate 107.2_cured_, and c) the 11,934 bp segment of pAQZ1 elimated through the curing experiments. Open reading frames (ORFs) shown in red depict resistance genes, ORFs in orange describe heavy metal efflux genes, ORFs in green represent transcriptional regulator genes, ORFs in blue depict recombinase genes, ORFs in purple describe replication genes, ORFs in white represent hypothetical proteins, and ORFs in yellow depict genes with other functions.

When pAQZ1 was compared to plasmid sequences deposited in GenBank, several regions of pAQZ1 showed high sequence similarity (>99%) with previously characterized *S. aureus* and coagulase-negative *Staphylococci* plasmids (CP017465 and CP023967). The complete sequence of pAQZ1, however, did not fully align with any single, previously characterized *S. aureus* or coagulase-negative *Staphylococci* plasmid. When queried against NCBI WGS, pAQZ1 showed high sequence similarity and a query coverage of 68% and 86%, respectively, to two previously sequenced contigs from two coagulase-negative *Staphylococcus* isolates (JZUM01000030.1 and QSTD01000014.1).

### Curing analysis in vitro confirms qacA4 is responsible for the elevated CHG MICs

Transformations of *S. epidermidis* TÜ1457 with pAQZ1 was attempted, but proved unsuccessful (data not shown). Thus, to confirm the observed association between *qacA4* and the elevated CHG MICs, we attempted to cure the *qacA4*-carrying, *smr*-negative *S. epidermidis* isolate 107.2 of the pAQZ1 plasmid. We took advantage of the ability of QacA to efflux ethidium bromide (18) to screen for colonies which lost *qacA4*. Cells without *qacA4* accumulate ethidium bromide in their cytoplasm and the resulting colonies fluoresce under UV radiation. Those retaining *qacA4* do not accumulate ethidium bromide and thus, the resulting colonies do not fluoresce.

After 11 successive passages in trypticase soy broth without selection, an isolate cured of *qacA4*, referred to as isolate 107.2_cured_, was identified. The CHG MIC of 107.2_cured_ was four-fold lower than that of 107.2 (Table 5). The 8-agent antimicrobial susceptibility profile of 107.2_cured_ was identical to that of the parental strain (Table 5). Sequencing of the 107.2_cured_ (Table S1) revealed recombination, presumably catalyzed by the recombinases on the plasmid, led to pAQZ1 eliminating an 11,934 bp segment and resulted in the formation of a new 17,497 bp plasmid. This new plasmid, pAQZ2 (MK046688; Figure 5b), retained the RepA protein of pAQZ1. The 11.9 kb segment lost in 107.2_cured_ not only contained *qacA4*, but also the *knt* kanamycin resistance gene, the *ble* bleomycin resistance gene, the partial β-lactamase operon, and several recombinases (Figure 5c). PCR testing further confirmed isolate 107.2_cured_ lost the 11.9 kb segment distinguishing pAQZ1 from pAQZ2 (data not shown). Isolate 107.2_cured_ contained one coding change in its chromosome when compared to the 107.2 parental strain. This coding change occurred in a GCN5-related N-acetyltransferase family protein (Gly225Glu).

**Table 5.**
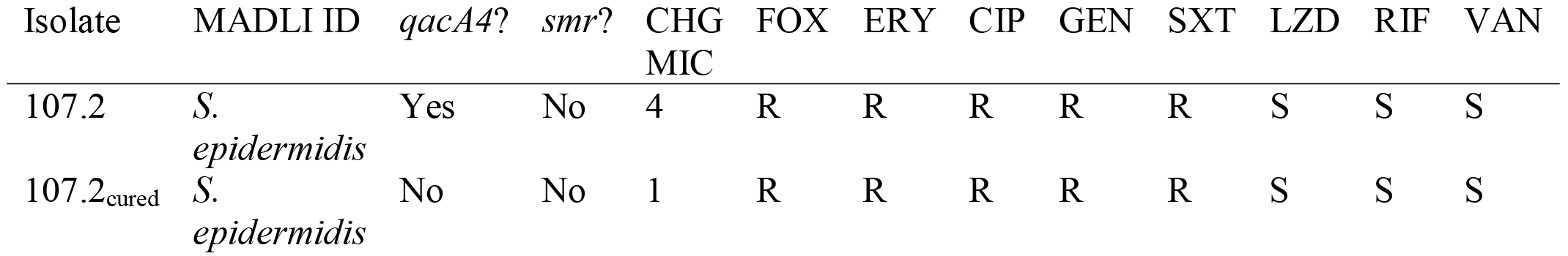
Comparison of the CHG resistance gene combinations and resistance patterns of isolates 107.2 and 107.2_cured_. The CHG MIC is measured in μg/mL. “R” denotes the isolate is resistant to the specified antimicrobial and “S” indicates the isolate is susceptible to the specified antimicrobial.

With the exception of *qacA4*, each of the genes present on the segment lost in isolate 107.2_cured_ was identified in at least one of the *qacA*-positive control isolates without elevated CHG MICs. One of these *qacA*-positive isolates, 110.3, contained all of these other 11 genes contained on the eliminated segment of pAQZ1.

### Isolates carrying qacA4 belong to the highly resistant and virulent S. epidermidis sequence type ST2

The isolates carrying *qacA4* harbored genes and mutations which confer resistance to several classes of commonly prescribed antimicrobials (Figure 6). Additionally, all of the sequenced *qacA4*-carrying isolates contained the biofilm formation operon, *icaADBC*. When classified by multilocus sequence typing (MLST), all these isolates belonged to the *S. epidermidis* sequence type ST2.

**Figure 6.**
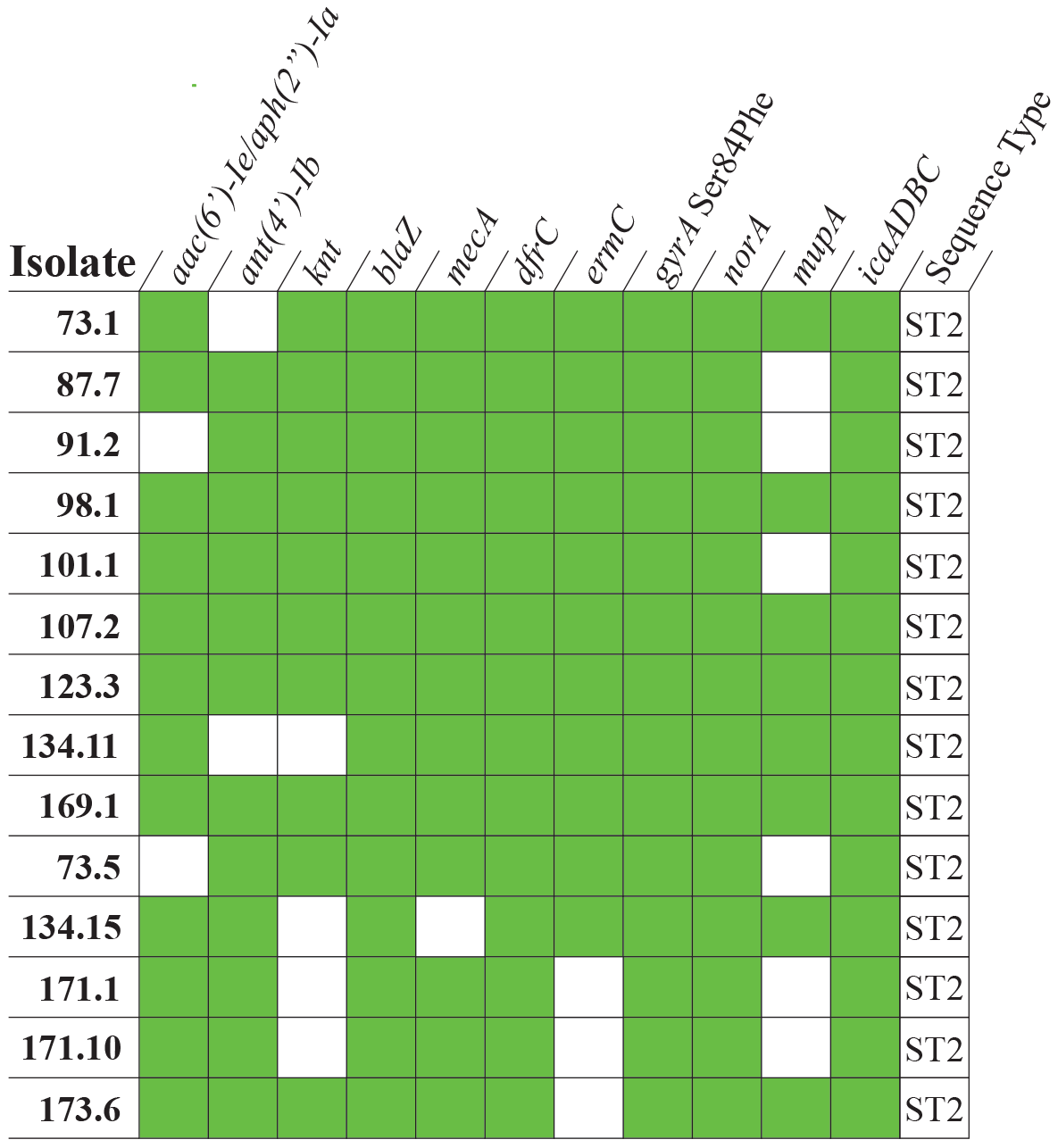
Presence-absence matrix displaying the antimicrobial resistance genes, resistance-associated mutations, and virulence genes identified in the isolates carrying *qacA4*. A green shaded box indicates the resistant or virulence determinant was identified in a given isolate. The sequence type of each isolate as determined by MLST is shown.

Five additional *qacAB*_*273*_-positive isolates with elevated CHG MICs, but displaying discordant susceptibility patterns (susceptible to methicillin, gentamicin, or erythromycin) were whole genome sequenced (Table S1). Each of the isolates carried *qacA4* and belonged to ST2. The divergent susceptibility patterns were explained by the absence of one or more resistance genes (Figure 6).

### qacA4-containing S. epidermidis isolates are distributed across North America

In total, 22 patients carried at least one cutaneous *S. epidermidis* isolate containing *qacA4* as confirmed by sequencing or as presumed through the isolate’s *qacAB*_*273*_-positive RFLP pattern and elevated CHG MIC. These 22 patients were enrolled at 14 study centers in 9 US states and 2 Canadian provinces (Figure 7). There was no obvious geographical clustering of the *qacA4*-carrying isolates.

**Figure 7.**
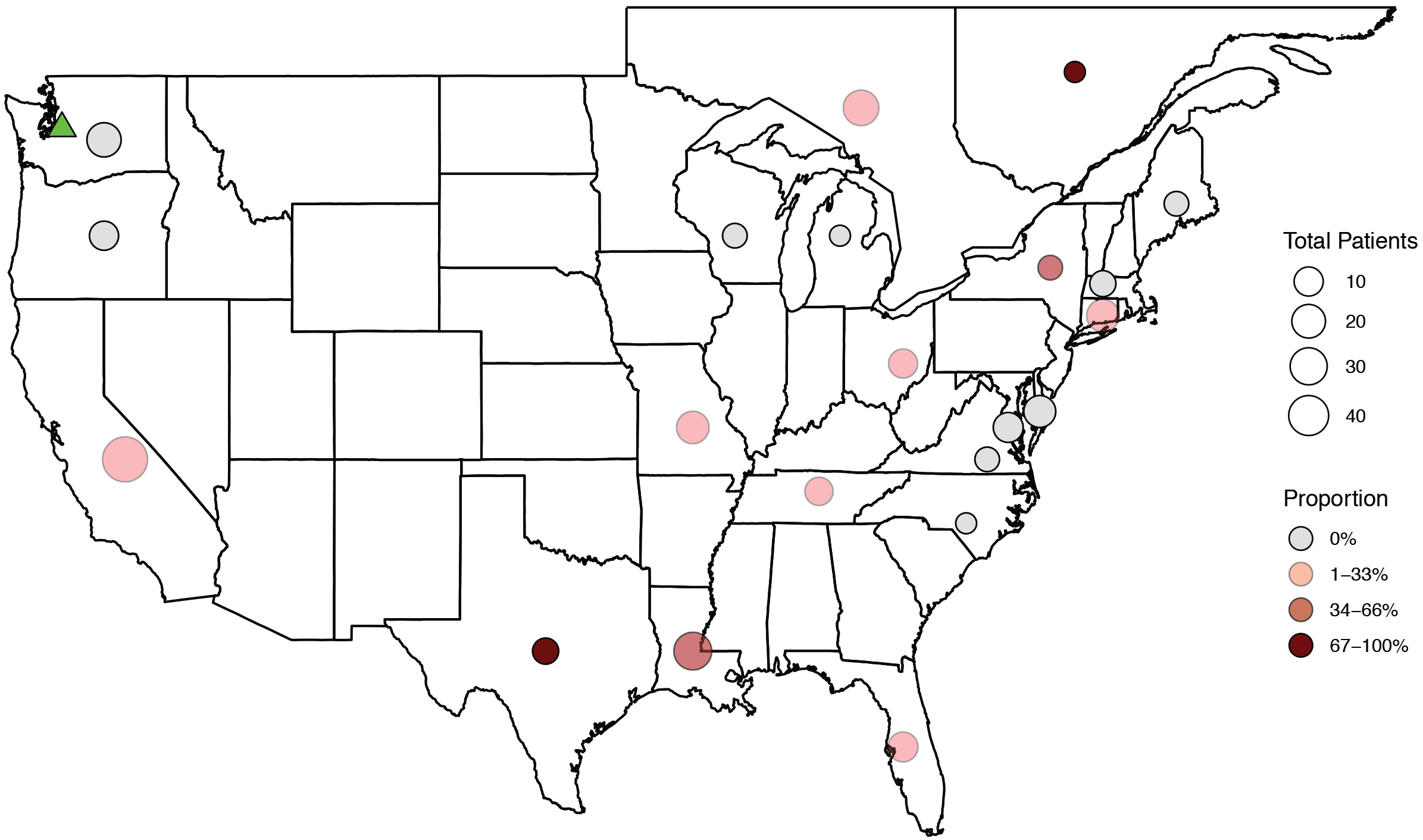
Geographic distribution of the patients with at least one *S. epidermidis* isolate carrying *qacA4*. The size of the circles represents the total number of patients enrolled in each state or province, and shade of the circle represents the proportion of the total patients with at least one *S. epidermidis* isolate. The green marker (Roach et al., 2015; Soma et al., 2012) represents isolates carrying *qacA4* collected outside of our study.

As shown in Figure 7, isolates containing *qacA4* existed outside of our study. A cutaneous *S. epidermidis* isolate containing *qacA4* (Table S1) was obtained from a CHG bathing pilot study (13) conducted at Seattle Children’s Hospital. Four clinical *S. epidermidis* isolates containing *qacA4* (SRA Accession Numbers: SRX761965, SRX762497, SRX762541, and SRX762777), identified through a query of the NCBI Sequence Read Archive, were obtained from a previous study (28) conducted at the University of Washington Medical Center. Susceptibility testing of these isolates revealed that all had an elevated CHG MIC (4 μg/mL).

## Discussion

In this study, we identified a novel *qacA* allele, termed *qacA4*, associated with an elevated CHG MIC in cutaneous *S. epidermidis* isolates and determined *qacA4* was contained on the novel pAQZ1 plasmid. We demonstrated *qacA4* was the determinant for the elevated CHG MIC by curing an isolate of the gene. Additionally, our analyses revealed isolates carrying *qacA4* displayed high rates of resistance to methicillin and other commonly prescribed antimicrobials, including erythromycin, ciprofloxacin, gentamicin, and sulfamethoxazole/trimethoprim. Whole genome sequencing revealed the isolates harbored several antimicrobial resistance determinants and resistance-associated mutations. Our analyses further demonstrated these isolates contained the chromosomally-encoded biofilm formation operon, *icaADBC* (2), and belonged to the highly resistant and pathogenetic ST2 clone (29). We identified isolates proven or presumed to carry *qacA4* from 22 patients enrolled in a multicenter, randomized controlled CHG bathing trial at 14 participating study centers across the United States and Canada. Furthermore, we identified *qacA4* in clinical *S. epidermidis* isolates collected in a prior study at a center without patients participating in our CHG bathing trial (28).

Previous studies have characterized three alleles of *qacA*: *qacA1*, *qacA2*, and *qacA3* (18, 21). Additional studies, however, have suggested clinical and environmental *Staphylococcus* isolates may carry novel alleles of *qacA* (30, 31). Our identification of multiple novel *qacA* alleles supports this suggestion that considerably more *qacA* allelic variation exists within human staphylococcal populations than previously appreciated.

The efflux pumps encoded by *qacA1*, *qacA2*, and *qacA3* do not exhibit any functional differences (21). However, the efflux potential of QacA has been shown to vary with its amino acid sequence (32–35). Despite this recognition, no studies have examined the functional differences associated with the sequence variation of *qacA* observed in clinical and environmental *Staphylococcus* isolates. As the CHG MIC of the *qacA4*-carrying isolates was significantly higher than the CHG MICs of the isolates carrying other alleles of *qacA*, our results suggest different alleles of *qacA* encode pumps with varying CHG efflux potentials. This is further supported by our identification of 4 novel *qacA* alleles from the nine *S. epidermidis* isolates with prototypical *qacA* restriction patterns and elevated CHG MICs. These novel alleles indicate other unique mutations may result in elevated CHG MICs and further underscore the importance of exploring the allelic variation of *qacA* in clinical and environmental *Staphylococcus* isolates.

It is tempting to speculate which of the two amino acid substitutions in QacA4, Ala157Gly and Ala378Val, is causal for the elevated CHG MIC observed in *qacA4*-carrying isolates. The Ala378Val substitution is particularly suspect as this mutation occurs in transmembrane segment 12. Transmembrane segment 12 is noteworthy when discussing the CHG efflux potential of QacA since a previous study demonstrated that this segment lines the CHG binding pocket (32). The Ala157Gly substitution was identified in an *S. epidermidis* isolate with the prototypical *qacA* RFLP pattern and an elevated CHG MIC. This may indicate the Ala157Gly substitution has a more causal role in the elevated CHG MICs. Future *in vitro* characterizations examining the structure-function relationship of the amino acid substitutions is merited.

Several studies have provided contradicting results as to whether the carriage of *qacA* influences the CHG MIC of *Staphylococcus* isolates (16, 19, 24, 36–40). These studies, however, did not distinguish between the *qacA* alleles carried by the isolates. Our findings highlight the importance of specifying the *qacA* allele carried by isolates when examining associations with CHG MICs: carriage of *qacA4*, as demonstrated by our curing analysis, results in a four-fold increase in the CHG MIC of an isolate, while carriage of other alleles may not increase the CHG MIC.

Screening for *qacA/B* has been used as a proxy for determining whether an isolate exhibits reduced susceptibility or tolerance to CHG (39, 41–43), typically defined as a CHG MIC ≥ 4 μg/mL (16, 39). All isolates carrying *qacA4* had a CHG MIC of 4 μg/mL, compared to just 10.0% of all *qacA/B*-positive isolates. Thus, screening for *qacA4*, rather than indiscriminately screening for *qacA/B*, may serve as a better indicator for reduced susceptibility to CHG in *Staphylococcus* spp.

Similar to previously described plasmids carrying *qacA/B* (22), pAQZ1 carries several genes involved in heavy metal efflux and a partial β-lactamase operon. As pAQZ1 only contains the β-lactamase transcriptional regulators, *blaI* and *blaR* (44), it is unclear if carriage of the plasmid influences β-lactam resistance. The kanamycin nucleotidyltransferase encoded by *knt* on pAQZ1 showed high sequence similarity to the kanamycin nucleotidyltransferase of *S. aureus* (X03408) and may contribute to aminoglycoside resistance (45). Since the cured isolate also contained other aminoglycoside resistant determinants, including *aac(6’)-Ie/aph(2”)-Ia* and *ant(4’)-Ib*, we were unable to assess the contribution of *knt* to aminoglycoside resistance.

All of the *qacA4*-carrying isolates we sequenced belonged to ST2, a *S. epidermidis* clone frequently implicated in device-associated infections (5, 29, 46–49). Consistent with previous studies (29, 46–48), our ST2 isolates contained genes and mutations which confer resistance to several classes of commonly prescribed antimicrobials. Additionally, our isolates contained genes associated with binding to foreign materials (5, 29), including the biofilm formation operon *icaADBC*. These results suggest *qacA4* may allow the highly resistant *S. epidermidis* ST2 clone to better persist following topical application of CHG and thus, further succeed as an opportunistic pathogen. However, as the concentration of CHG used in clinical settings (8, 10) is much higher than tested *in vitro* (2,000 μg/mL versus 4 μg/mL), further study is required to fully understand the clinical implications of carriage of *qacA4* by the ST2 clone.

Our results suggest *qacA4* is distributed in pediatric oncology populations at centers across the United States and Canada. Four isolates carrying *qacA4* were also identified from a prior study conducted at an institution without patients participating in our CHG bathing trial (28). These four isolates were collected from patients in intensive care units where CHG bathing was standard of care (Estella Whimbey, personal communication). With both the *S. epidermidis* ST2 clone and *qacA* widely disseminated throughout healthcare settings globally (29, 39), broader screening may reveal *qacA4* follows this wide global distribution.

Our study was limited by the nature of the RFLP screening analysis. While our method of screening for *qacA/B* allowed us to identify all the isolates with a mutation at positions 1131 to 1134, we were unable to easily detect the other novel *qacA* alleles that may have been present in our study population. From just the 35 *qacA*-positive and *qacAB*_*273*_-positive isolates we sequenced, we identified 11 novel *qacA* alleles, and, as we demonstrated, at least one of these alleles exhibits functional difference with respect to CHG efflux. This emphasizes the necessity of using sequence to screen for allelic variation in resistance determinants, especially in those determinants in which allelic variation has been underappreciated. Furthermore, reflecting the difficulty of performing transformations in *Staphylococcus* spp. (50), we were unable to perform a gain-of-function analysis for *qacA4* despite trying three different methods of preparing electrocompetent cells and two separate electroporation conditions for each cell preparation. Despite this limitation, we were able to perform a loss-of-function analysis to confirm the role of *qacA4* in the elevated CHG MIC. It is remarkable that the loss-of-function was achieved by recombination and that the cured isolate retained more than half of pAQZ1. Beyond *qacA4*, each of the 11 other genes contained on the segment lost in the cured isolate may explain the four-fold decrease in the CHG MIC exhibited by this cured isolate. Many of these genes, however, have well-described functions unrelated to CHG efflux (44, 45, 51–53). Additionally, we identified each of the other 11 genes in an isolate without an elevated CHG MIC. Thus, the decrease in the CHG MIC observed in the cured isolate is most consistent with the loss of *qacA4*.

Our results highlight the importance of screening for allelic variation in *qacA*. Just as a single SNV between *qacA* and *qacB* accounts for the differing substrate specificities of the resulting efflux pumps (18), the three SNVs of *qacA4* are associated with a four-fold increase in the CHG MIC. Further study should focus on understanding the functional differences of the various *qacA* alleles identified in clinical and environmental *Staphylococcus* isolates. Moreover, our results indicate the highly resistant *S. epidermidis* ST2 clone (29) carries *qacA4*. Future study is required to understand if frequent usage of CHG selects for *qacA4* and this pathogenic clone of *S. epidermidis*.

## Material and Methods

### Collection and identification of cutaneous *Staphylococcus* isolates

Skins swabs were obtained from patients between 2 months and 21 years of age undergoing allogeneic hematopoietic cell transplantation or treatment for cancer who were enrolled in a randomized double-blind placebo-controlled trial of CHG bathing versus control bathing conducted at 37 centers in the United States and Canada from January 2014 to April 2017 (Children’s Oncology Group ACCL1034). The study was approved by the National Cancer Institute’s Pediatric Central Institution Review Board as well as the local review boards at participating institutions, if required.

Samples were obtained by swabbing a 3×3 cm area on the side or back of the neck and axilla regions with a sterile nylon swab (Copan Diagnostics) for 20 seconds and transported in 1 mL of the accompanying liquid Amies medium. The swab and Amies medium were vigorously vortexed and the medium was plated on the following agar plates: Tryptic Soy with 5% Sheep’s Blood (Remel), Chocolate (Remel), Sabouraud Dextrose (Remel), MacConkey (Remel), and Mannitol Salt (Remel). The plates were incubated at 35°C for 48 hours. *Staphylococcus* isolates were identified via matrix-assisted laser desorption/ionization time of flight mass spectrometry (MALDI-TOF MS).

Isolates were prepared for MALDI-TOF MS according to the manufacturer’s Direct Transfer Sample Preparation procedure (54). A MicroFlex LT mass spectrometer (Bruker Daltonics, Inc) operated in the positive linear mode with FlexControl software (version 3.4, Bruker) was used to obtain spectra. The resulting spectra were processed and classified using Biotyper software (version 3.2, Bruker). Identification results were interpreted according to the manufacturer’s guidelines. The isolates, and the corresponding phenotypic information, included in the study are presented in the Dataset S1.

Five additional *qacA4*-carrying isolates were obtained for phenotypic testing: one was collected during a pilot study conducted at Seattle Children’s Hospital (13), and the other four were collected in a previous study at the University of Washington Medical Center (28).

### Antimicrobial susceptibility testing

Following CLSI guidelines (55), susceptibility testing was performed by disk diffusion (Benton, Dickinson, and Company) for the following antimicrobials: ERY (15 μg), CIP (5 μg), GEN (10 μg), LZD (30 μg), cefoxitin (FOX; 30 μg), RIF (5 μg), and SXT (23.75/1.25 μg). Additionally, VAN (0.016 – 256 μg/mL) susceptibility testing was performed using the Etest (BioMérieux) MIC method. FOX was used as a surrogate for methicillin susceptibility per CLSI guidelines (55). Isolates were classified as resistant, intermediate, or susceptible to a given agent using the breakpoints specified by CLSI (56).

CHG MICs (0.0625 – 64 μg/mL) were determined via the broth microdilution method (56, 57). For the CHG MICs, the following strains were included as controls: *Pseudomonas aeruginosa* ATCC 27853, *Staphylococcus aureus* ATCC 25923, *Staphylococcus epidermidis* (Laboratory control strain) (13), *Escherichia coli* ATCC 25922.

### Detection of *qacA/B* and *smr*

Isolates were tested for the presence of *qacA/B* via PCR using AmpliTaq DNA Polymerase (Applied Biosystems) following the manufacturer’s instruction for the reaction mixture (58) with the previously described primers and reaction conditions (21). To distinguish between *qacA* and *qacB*, the PCR products were digested with *AluI* and the resulting fragments were visualized with agarose gel electrophoresis (21). Control *qacA*-positive and *qacB*-positive strains were obtained from Dr. Nobumichi Kobayashi.

Isolates were additionally screened for the presence of *smr* via PCR with the following primers: F – 5’AAAACAATGCAACACCTACCAC3’ and R – 5’ATGCGATGTTCCGAAAATGT3’. The following reactions conditions were used: an initial denaturation for 3 minutes at 95°C, followed by 30 cycles of denaturation for 1 minute at 95°C, primer annealing for 30 seconds at 55°C, and elongation for 1 minute at 72°C, and completed with a final elongation for 10 minutes at 72°C. A control *smr*-positive strain was obtained from Dr. Arnold Bayer.

### Statistical testing

Kruskal-Wallis or Wilcoxon rank-sum tests were used to assess differences in the distribution of CHG MICs between *qac* groups. Fisher’s exact test was used to assess the proportion of isolates resistant to commonly used antimicrobials by *qac* group. All analyses were first performed on all isolates and then repeated using one randomly chosen isolate per patient per *qac* group. Analyses were performed using STATA version 14 (College Station, TX) and R version 3.3.2 (R Core Team).

### Whole genome Sequencing

In total, the genomes of 40 *Staphylococcus* spp. isolates were sequenced. An initial group of 10 *qacA*-positive, 4 *qacB*-positive, and 10 *qacAB*_*273*_-positive isolates was selected for whole genome sequencing. The pool of *qacA*-positive and *qacB*-positive isolates was restricted to those identified as *S. epidermidis* since all of the *qacAB*_*273*_-positive with an elevated CHG MIC were identified as *S. epidermidis*. The pool of isolates was then further restricted to the first *qacA*-positive isolate from 10 patients randomly chosen from the 91 patients with at least one *qacA*-positive isolate, the first *qacAB*_*273*_-positive isolate obtained from 10 patients randomly chosen from the 24 patients with at least one *qacAB*_*273*_-positive isolate, and the first *qacB*-positive isolate from all 4 patients with at least one *qacB*-positive isolate. One of the initially selected *qacAB*_*273*_-positive isolates contained two alleles of *qacA*, with one being *qacA4*. As a result, this isolate was excluded from all subsequent analyses. Additional isolates were selected for whole genome sequencing based on their antimicrobial susceptibility phenotypes: 2 *qacAB*_*273*_-positive isolates were chosen as they did not have elevated CHG MICs, 9 *qacA*-positive isolates were selected as they had elevated CHG MICs, and 5 *qacAB*_*273*_-positive isolates were chosen as they had discordant antimicrobial susceptibility patterns (susceptible to FOX, GEN, or ERY).

Isolates were grown in Brain Heart Infusion broth (Remel) for 24 hours at 37°C at a constant shaking of 150 rpm. DNA was extracted from these isolates using the QIAamp DNA Mini Kit (Qiagen) while following the manufacturer’s protocol for isolating DNA from Gram-positive bacteria (59). An initial cell lysis step was completed using a 200 μg/mL lysostaphin solution (Sigma-Aldrich).

Libraries were prepared with the KAPA Hyper Prep Kit. Isolates were sequenced to at least 27X coverage using 2×300 bp Illumina MiSeq runs. De novo assemblies were constructed with SPAdes, annotated via prokka, and visualized using Geneious v.10.2.3.

Multilocus sequence typing was completed by uploading the assemblies to PubMLST’s *Staphylococcus epidermidis* MLST website (https://pubmlst.org/sepidermidis/) (60). Resistance genes, limited to only those matching “Perfect” and “Strict” criteria, were detected with the Comprehensive Antibiotic Resistance Database’s Resistance Gene Identifier (https://card.mcmaster.ca/analyze/rgi) (61). The sequence of the DNA gyrase A protein in our isolates was compared with the DNA gyrase A protein of *S. epidermidis* ATCC12228 (AE015929), a ciprofloxacin-sensitive strain, to determine the identity of the residue at position 84. Isolates containing the Ser84Phe mutation were determined to contain a *gyrA* gene conferring resistance to fluoroquinolones (62). The assemblies were additionally screened for virulence genes including *icaADBC* (U43366), *aap* (KJ920749), and *bhp* (AY028618).

### Curing of *qacA4*

An isolate carrying *qacA4*, 107.2, was selected for the curing analysis as this isolate did not contain *smr*, which can efflux ethidium bromide. The isolate was successively passaged in Tryptic Soy broth (Remel) in four separate curing conditions: No Selection – incubated for 24 hours at 37°C at a constant shaking of 150 rpm; Increased Temperature – incubated for 24 hours at 42°C; Increasing Sub-Inhibitory Concentrations of Sodium Dodecyl Sulfate (Sigma-Aldrich) – incubated for 24 hours at 37°C at a constant shaking of 150 rpm with 0.001% to 0.01% sodium dodecyl sulfate; and Increasing Sub-Inhibitory Concentrations of Novobiocin (Sigma-Aldrich) – incubated for 24 hours at 37°C at a constant shaking of 150 rpm with 0.01 μg/mL to 0.1 μg/mL novobiocin.

After each passage, broths were plated onto Tryptic Soy agar plates (Remel) containing 0.375 μg/mL of filter-sterilized ethidium bromide (VWR). The plates were incubated at 35°C for 48 hours. Screening for cured strains was completed with ultraviolet light as previously described (37). PCR was used to confirm the cured strain eliminated *qacA4*. Whole genome sequencing was used to confirm the cured strain contained minimal chromosomal mutations as compared to the parental strain.

Three PCR reactions were conducted on the plasmids predicted from whole genome sequencing of the isolates. To confirm the circular nature of the contig presumed to be a plasmid, PCR was conducted with the following primers: F – 5’GGCTACTGTTGTTTTACCTACACCACC3’ and R – 5’GCATACATAACCTTTGCGTCAGTTGTC3’. To confirm curing resulted in the formation of a novel plasmid, PCR was conducted with the following primers: F – 5’CCATTGTGGCGTCATTTCACGGC3’ and R – 5’CGGCGAAATCCTTGAGCCATATCTG3’ and F – 5’GAAGAATCTGTAGTGGGCGCTG3’ and R – 5’GATGAAAGTTGCTACTAGTGCTCC3’. The following reactions conditions were used: an initial denaturation for 3 minutes at 95°C, followed by 30 cycles of denaturation for 1 minute at 95°C, primer annealing for 30 seconds at 53°C, 57°C, or 52°C, respectively, and elongation for 1 minute at 72°C, and completed with a final elongation for 10 minutes at 72°C.

### Transformation of pAQZ1 into *S. epidermidis* TÜ1457

In preparation for the extraction of the pAQZ1 plasmid, the *qacA4*-carrying *S. epidermidis* isolate 107.2 was grown Tryptic Soy Broth (Remel) for 24 hours at 37°C at a constant shaking of 150 rpm. The plasmid was extracted using the QIAprep Spin Miniprep Kit (Qiagen) following the manufacturer’s instructions (63).

The pan-susceptible *S. epidermidis* TÜ1457 strain (64) was used for the transformations. Three previously described methods were used for preparing electrocompetent cells (50, 65, 66). For electroporation, 100 μL of the prepared cells were mixed with 100 ng of pAQZ1 DNA in a 1-mm electroporation cuvette (Bio-Rad). Two electroporation conditions were used for each preparation of electrocompetent cells: 21 kV/cm, 100 Ω, and 25 μF; and 23 kV/cm, 100 Ω, and 25 μF. The pulsed cells were resuspended 1000 μL of broth, with the type of broth being selected based on the previously described methods, and incubated at 37°C at a constant shaking for 150 rpm for 1 hour. The cells were plated onto Tryptic Soy Agar plates (Remel) containing either 2 μg/mL CHG (Sigma-Aldrich), 15 μg/mL ethidium bromide (VWR), or 10 μg/mL of kanamycin (Sigma-Aldrich) and incubated overnight at 37°C.

### Accession Numbers

The sequence of *qacA4* was deposited in GenBank under accession number MK040360. The accession numbers for the additional 10 novel *qacA* alleles identified in this study are listed in Table S2. The sequences of pAQZ1 and pAQZ2 were deposited under accession numbers MK046687 and MK046688, respectively. Draft genome assemblies are available in GenBank under the study accession PRJNA415995. The accession numbers, as well as the phenotypic data, for the individual isolates sequenced are displayed in Table S1.

## Supporting information

## Acknowledgments

The authors would like to thank the patients who agreed to participate in this study as well as the research teams at all the involved sites and the ACCL1034 committee. The authors would also like to thank Dr. Stephen Salipante for sharing previously collected bacterial isolates, Dr. Paul Fey for providing the *S. epidermidis* TÜ1457 strain, Dr. Nobumichi Kobayashi for supplying the *qacA/B*-positive control strains, and Dr. Arnold Bayer for sharing the *smr*-positive control strain.

This work was supported by the National Cancer Institute at the National Institutes of Health [R01CA163394 and UG1CA189955]. The content is solely the responsibility of the authors and does not necessarily represent the official views of the National Institutes of Health.

The research was also supported by St. Baldrick’s Foundation.

## References

1. Becker K, Heilmann C, Peters G. 2014. Coagulase-Negative Staphylococci. Clinical Microbiology Reviews 27:870–926.

2. Gerke C, Kraft A, Süssmuth R, Schweitzer O, Götz F. 1998. Characterization of the N-acetylglucosaminyltransferase activity involved in the biosynthesis of the Staphylococcus epidermidis polysaccharide intercellular adhesin. J Biol Chem 273:18586–18593.

3. Fey PD, Olson ME. 2010. Current concepts in biofilm formation of Staphylococcus epidermidis. Future Microbiology 5:917–933.

4. Hussain M, Herrmann M, von Eiff C, Perdreau-Remington F, Peters G. 1997. A 140-kilodalton extracellular protein is essential for the accumulation of Staphylococcus epidermidis strains on surfaces. Infect Immun 65:519–524.

5. Otto M. 2009. Staphylococcus epidermidis — the “accidental” pathogen. Nature Reviews Microbiology 7:555–567.

6. Diekema DJ, Pfaller MA, Schmitz FJ, Smayevsky J, Bell J, Jones RN, Beach M, SENTRY Participants Group. 2001. Survey of Infections Due to Staphylococcus Species: Frequency of Occurrence and Antimicrobial Susceptibility of Isolates Collected in the United States, Canada, Latin America, Europe, and the Western Pacific Region for the SENTRY Antimicrobial Surveillance Program, 1997–1999. Clinical Infectious Diseases 32:S114–S132.

7. Edmiston CE, Bruden B, Rucinski MC, Henen C, Graham MB, Lewis BL. 2013. Reducing the risk of surgical site infections: Does chlorhexidine gluconate provide a risk reduction benefit? American Journal of Infection Control 41:S49–S55.

8. Milstone AM, Elward A, Song X, Zerr DM, Orscheln R, Speck K, Obeng D, Reich NG, Coffin SE, Perl TM. 2013. Daily chlorhexidine bathing to reduce bacteraemia in critically ill children: a multicentre, cluster-randomised, crossover trial. The Lancet 381:1099–1106.

9. Bleasdale SC, Trick WE, Gonzalez IM, Lyles RD, Hayden MK, Weinstein RA. 2007. Effectiveness of Chlorhexidine Bathing to Reduce Catheter-Associated Bloodstream Infections in Medical Intensive Care Unit Patients. Archives of Internal Medicine 167:2073.

10. Climo MW, Yokoe DS, Warren DK, Perl TM, Bolon M, Herwaldt LA, Weinstein RA, Sepkowitz KA, Jernigan JA, Sanogo K, Wong ES. 2013. Effect of Daily Chlorhexidine Bathing on Hospital-Acquired Infection. New England Journal of Medicine 368:533–542.

11. Mimoz O, Karim A, Mercat A, Cosseron M, Falissard B, Parker F, Richard C, Samii K, Nordmann P. 1999. Chlorhexidine compared with povidone-iodine as skin preparation before blood culture. A randomized, controlled trial. Ann Intern Med 131:834–837.

12. Marlowe L, Mistry RD, Coffin S, Leckerman KH, McGowan KL, Dai D, Bell LM, Zaoutis T. 2010. Blood Culture Contamination Rates after Skin Antisepsis with Chlorhexidine Gluconate versus Povidone-Iodine in a Pediatric Emergency Department. Infection Control & Hospital Epidemiology 31:171–176.

13. Soma V, Qin X, Zhou C, Adler A, Berry J, Zerr D. 2012. The effects of daily chlorhexidine bathing on cutaneous bacterial isolates: a pilot study. Infection and Drug Resistance 75–78.

14. Vernon MO, Hayden MK, Trick WE, Hayes RA, Blom DW, Weinstein RA. 2006. Chlorhexidine Gluconate to Cleanse Patients in a Medical Intensive Care Unit: The Effectiveness of Source Control to Reduce the Bioburden of Vancomycin-Resistant Enterococci. Archives of Internal Medicine 166:306.

15. Block C, Furman M. 2002. Association between intensity of chlorhexidine use and micro-organisms of reduced susceptibility in a hospital environment. J Hosp Infect 51:201–206.

16. Wang J-T, Sheng W-H, Wang J-L, Chen D, Chen M-L, Chen Y-C, Chang S-C. 2008. Longitudinal analysis of chlorhexidine susceptibilities of nosocomial methicillin-resistant Staphylococcus aureus isolates at a teaching hospital in Taiwan. Journal of Antimicrobial Chemotherapy 62:514–517.

17. Vali L, Davies SE, Lai LLG, Dave J, Amyes SGB. 2008. Frequency of biocide resistance genes, antibiotic resistance and the effect of chlorhexidine exposure on clinical methicillin-resistant Staphylococcus aureus isolates. Journal of Antimicrobial Chemotherapy 61:524–532.

18. Paulsen IT, Brown MH, Littlejohn TG, Mitchell BA, Skurray RA. 1996. Multidrug resistance proteins QacA and QacB from Staphylococcus aureus: membrane topology and identification of residues involved in substrate specificity. Proceedings of the National Academy of Sciences 93:3630–3635.

19. Mitchell BA, Brown MH, Skurray RA. 1998. QacA multidrug efflux pump from Staphylococcus aureus: comparative analysis of resistance to diamidines, biguanidines, and guanylhydrazones. Antimicrob Agents Chemother 42:475–477.

20. Brown MH, Skurray RA. 2001. Staphylococcal multidrug efflux protein QacA. J Mol Microbiol Biotechnol 3:163–170.

21. Alam MM, Kobayashi N, Uehara N, Watanabe N. 2003. Analysis on Distribution and Genomic Diversity of High-Level Antiseptic Resistance Genes qacA and qacB in Human Clinical Isolates of Staphylococcus aureus. Microbial Drug Resistance 9:109–121.

22. Leelaporn A, Paulsen IT, Tennent JM, Littlejohn TG, Skurray RA. 1994. Multidrug resistance to antiseptics and disinfectants in coagulase-negative staphylococci. Journal of Medical Microbiology 40:214–220.

23. Paulsen IT, Brown MH, Dunstan SJ, Skurray RA. 1995. Molecular characterization of the staphylococcal multidrug resistance export protein QacC. Journal of Bacteriology 177:2827–2833.

24. Littlejohn TG, Paulsen IT, Gillespie MT, Tennent JM, Midgley M, Jones IG, Purewal AS, Skurray RA. 1992. Substrate specificity and energetics of antiseptic and disinfectant resistance in Staphylococcus aureus. FEMS Microbiol Lett 74:259–265.

25. Bjorland J, Sunde M, Waage S. 2001. Plasmid-Borne smr Gene Causes Resistance to Quaternary Ammonium Compounds in Bovine Staphylococcus aureus. Journal of Clinical Microbiology 39:3999–4004.

26. Weaver KE, Kwong SM, Firth N, Francia MV. 2009. The RepA_N replicons of Gram-positive bacteria: A family of broadly distributed but narrow host range plasmids. Plasmid 61:94–109.

27. Kwong SM, Ramsay JP, Jensen SO, Firth N. 2017. Replication of Staphylococcal Resistance Plasmids. Frontiers in Microbiology 8.

28. Roach DJ, Burton JN, Lee C, Stackhouse B, Butler-Wu SM, Cookson BT, Shendure J, Salipante SJ. 2015. A Year of Infection in the Intensive Care Unit: Prospective Whole Genome Sequencing of Bacterial Clinical Isolates Reveals Cryptic Transmissions and Novel Microbiota. PLOS Genetics 11:e1005413.

29. Schoenfelder SMK, Lange C, Eckart M, Hennig S, Kozytska S, Ziebuhr W. 2010. Success through diversity – How Staphylococcus epidermidis establishes as a nosocomial pathogen. International Journal of Medical Microbiology 300:380–386.

30. Wassenaar T, Ussery D, Nielsen L, Ingmer H. 2015. Review and phylogenetic analysis of qac genes that reduce susceptibility to quaternary ammonium compounds in Staphylococcus species. European Journal of Microbiology and Immunology 5:44–61.

31. Hijazi K, Mukhopadhya I, Abbott F, Milne K, Al-Jabri ZJ, Oggioni MR, Gould IM. 2016. Susceptibility to chlorhexidine amongst multidrug-resistant clinical isolates of Staphylococcus epidermidis from bloodstream infections. International Journal of Antimicrobial Agents 48:86–90.

32. Hassan KA, Skurray RA, Brown MH. 2007. Transmembrane Helix 12 of the Staphylococcus aureus Multidrug Transporter QacA Lines the Bivalent Cationic Drug Binding Pocket. Journal of Bacteriology 189:9131–9134.

33. Wu J, Hassan KA, Skurray RA, Brown MH. 2008. Functional analyses reveal an important role for tyrosine residues in the staphylococcal multidrug efflux protein QacA. BMC Microbiology 8:147.

34. Hassan KA, Galea M, Wu J, Mitchell BA, Skurray RA, Brown MH. 2006. Functional effects of intramembranous proline substitutions in the staphylococcal multidrug transporter QacA. FEMS Microbiology Letters 263:76–85.

35. Xu Z, O’Rourke BA, Skurray RA, Brown MH. 2006. Role of Transmembrane Segment 10 in Efflux Mediated by the Staphylococcal Multidrug Transport Protein QacA. Journal of Biological Chemistry 281:792–799.

36. Sheng W-H, Wang J-T, Lauderdale T-L, Weng C-M, Chen D, Chang S-C. 2009. Epidemiology and susceptibilities of methicillin-resistant Staphylococcus aureus in Taiwan: emphasis on chlorhexidine susceptibility. Diagnostic Microbiology and Infectious Disease 63:309–313.

37. Costa SS, Ntokou E, Martins A, Viveiros M, Pournaras S, Couto I, Amaral L. 2010. Identification of the plasmid-encoded qacA efflux pump gene in meticillin-resistant Staphylococcus aureus (MRSA) strain HPV107, a representative of the MRSA Iberian clone. International Journal of Antimicrobial Agents 36:557–561.

38. Skovgaard S, Larsen MH, Nielsen LN, Skov RL, Wong C, Westh H, Ingmer H. 2013. Recently introduced qacA/B genes in Staphylococcus epidermidis do not increase chlorhexidine MIC/MBC. Journal of Antimicrobial Chemotherapy.

39. Horner C, Mawer D, Wilcox M. 2012. Reduced susceptibility to chlorhexidine in staphylococci: is it increasing and does it matter? Journal of Antimicrobial Chemotherapy 67:2547–2559.

40. Hayden MK, Lolans K, Haffenreffer K, Avery TR, Kleinman K, Li H, Kaganov RE, Lankiewicz J, Moody J, Septimus E, Weinstein RA, Hickok J, Jernigan J, Perlin JB, Platt R, Huang SS. 2016. Chlorhexidine and Mupirocin Susceptibility of Methicillin-Resistant Staphylococcus aureus Isolates in the REDUCE-MRSA Trial. Journal of Clinical Microbiology 54:2735–2742.

41. McClure J-A, Zaal DeLongchamp J, Conly JM, Zhang K. 2017. Novel Multiplex PCR Assay for Detection of Chlorhexidine-Quaternary Ammonium, Mupirocin, and Methicillin Resistance Genes, with Simultaneous Discrimination of Staphylococcus aureus from Coagulase-Negative Staphylococci. Journal of Clinical Microbiology 55:1857–1864.

42. Noguchi N. 2005. Susceptibilities to antiseptic agents and distribution of antiseptic-resistance genes qacA/B and smr of methicillin-resistant Staphylococcus aureus isolated in Asia during 1998 and 1999. Journal of Medical Microbiology 54:557–565.

43. Warren DK, Prager M, Munigala S, Wallace MA, Kennedy CR, Bommarito KM, Mazuski JE, Burnham C-AD. 2016. Prevalence of qacA/B Genes and Mupirocin Resistance Among Methicillin-Resistant Staphylococcus aureus (MRSA) Isolates in the Setting of Chlorhexidine Bathing Without Mupirocin. Infection Control & Hospital Epidemiology 37:590–597.

44. Hackbarth CJ, Chambers HF. 1993. blaI and blaR1 regulate beta-lactamase and PBP 2a production in methicillin-resistant Staphylococcus aureus. Antimicrob Agents Chemother 37:1144–1149.

45. Pedersen LC, Benning MM, Holden HM. 1995. Structural investigation of the antibiotic and ATP-binding sites in kanamycin nucleotidyltransferase. Biochemistry 34:13305–13311.

46. Li M, Wang X, Gao Q, Lu Y. 2009. Molecular characterization of Staphylococcus epidermidis strains isolated from a teaching hospital in Shanghai, China. Journal of Medical Microbiology 58:456–461.

47. Iorio NLP, Caboclo RF, Azevedo MB, Barcellos AG, Neves FPG, Domingues RMCP, dos Santos KRN. 2012. Characteristics related to antimicrobial resistance and biofilm formation of widespread methicillin-resistant Staphylococcus epidermidis ST2 and ST23 lineages in Rio de Janeiro hospitals, Brazil. Diagnostic Microbiology and Infectious Disease 72:32–40.

48. Hellmark B, Söderquist B, Unemo M, Nilsdotter-Augustinsson Å. 2013. Comparison of Staphylococcus epidermidis isolated from prosthetic joint infections and commensal isolates in regard to antibiotic susceptibility, agr type, biofilm production, and epidemiology. International Journal of Medical Microbiology 303:32–39.

49. Widerström M, McCullough CA, Coombs GW, Monsen T, Christiansen KJ. 2012. A Multidrug-Resistant Staphylococcus epidermidis Clone (ST2) Is an Ongoing Cause of Hospital-Acquired Infection in a Western Australian Hospital. Journal of Clinical Microbiology 50:2147–2151.

50. Monk IR, Shah IM, Xu M, Tan M-W, Foster TJ. 2012. Transforming the Untransformable: Application of Direct Transformation To Manipulate Genetically Staphylococcus aureus and Staphylococcus epidermidis. mBio 3.

51. Grkovic S, Brown MH, Roberts NJ, Paulsen IT, Skurray RA. 1998. QacR is a repressor protein that regulates expression of the Staphylococcus aureus multidrug efflux pump QacA. J Biol Chem 273:18665–18673.

52. Novick RP. 1989. Staphylococcal Plasmids and their Replication. Annual Review of Microbiology 43:537–563.

53. Sugiyama M, Kumagai T, Matsuo H, Bhuiyan MZ, Ueda K, Mochizuki H, Nakamura N, Davies JE. 1995. Overproduction of the bleomycin-binding proteins from bleomycin-producing Streptomyces verticillus and a methicillin-resistant Staphylococcus aureus in Escherichia coli and their immunological characterisation. FEBS Lett 362:80–84.

54. Bruker Daltonics. 2015. Instructions for Use Bruker Matrix HCCA.

55. CLSI. 2018. Performance Standards for Antimicrobial Disk Susceptibility Tests (M100).

56. CLSI. 2012. Methods for Dilution Antimicrobial Susceptibility Tests for Bacteria that Grow Aerobically (M07-A9).

57. CLSI. 1999. Methods for Determining Bactericidal Activity of Antimicrobial Agents (M26-A).

58. Applied Biosystems. 2014. AmpliTaq DNA Polymerase Protocol.

59. QIAGEN. 2016. QIAamp^®^ DNA Mini and Blood Mini Handbook.

60. Jolley KA, Bray JE, Maiden MCJ. 2018. Open-access bacterial population genomics: BIGSdb software, the PubMLST.org website and their applications. Wellcome Open Research 3:124.

61. Jia B, Raphenya AR, Alcock B, Waglechner N, Guo P, Tsang KK, Lago BA, Dave BM, Pereira S, Sharma AN, Doshi S, Courtot M, Lo R, Williams LE, Frye JG, Elsayegh T, Sardar D, Westman EL, Pawlowski AC, Johnson TA, Brinkman FSL, Wright GD, McArthur AG. 2017. CARD 2017: expansion and model-centric curation of the comprehensive antibiotic resistance database. Nucleic Acids Research 45:D566–D573.

62. Sreedharan S, Peterson LR, Fisher LM. 1991. Ciprofloxacin resistance in coagulase-positive and -negative staphylococci: role of mutations at serine 84 in the DNA gyrase A protein of Staphylococcus aureus and Staphylococcus epidermidis. Antimicrob Agents Chemother 35:2151–2154.

63. QIAGEN. 2015. QIAprep^®^ Spin Miniprep Kit.

64. Galac MR, Stam J, Maybank R, Hinkle M, Mack D, Rohde H, Roth AL, Fey PD. 2017. Complete Genome Sequence of Staphylococcus epidermidis 1457. Genome Announcements 5.

65. Andreote FD, Gullo MJM, de Souza Lima AO, Júnior WM, Azevedo JL, Araújo WL. 2004. Impact of genetically modified Enterobacter cloacae on indigenous endophytic community of Citrus sinensis seedlings. J Microbiol 42:169–173.

66. Schenk S, Laddaga RA. 1992. Improved method for electroporation of Staphylococcus aureus. FEMS Microbiol Lett 73:133–138.

