## Supplementary material for "A novel *qacA* allele results in an elevated chlorhexidine glu conate minimum inhibitory concentration in cutaneous *Staphylococcus epidermidis* isolates"

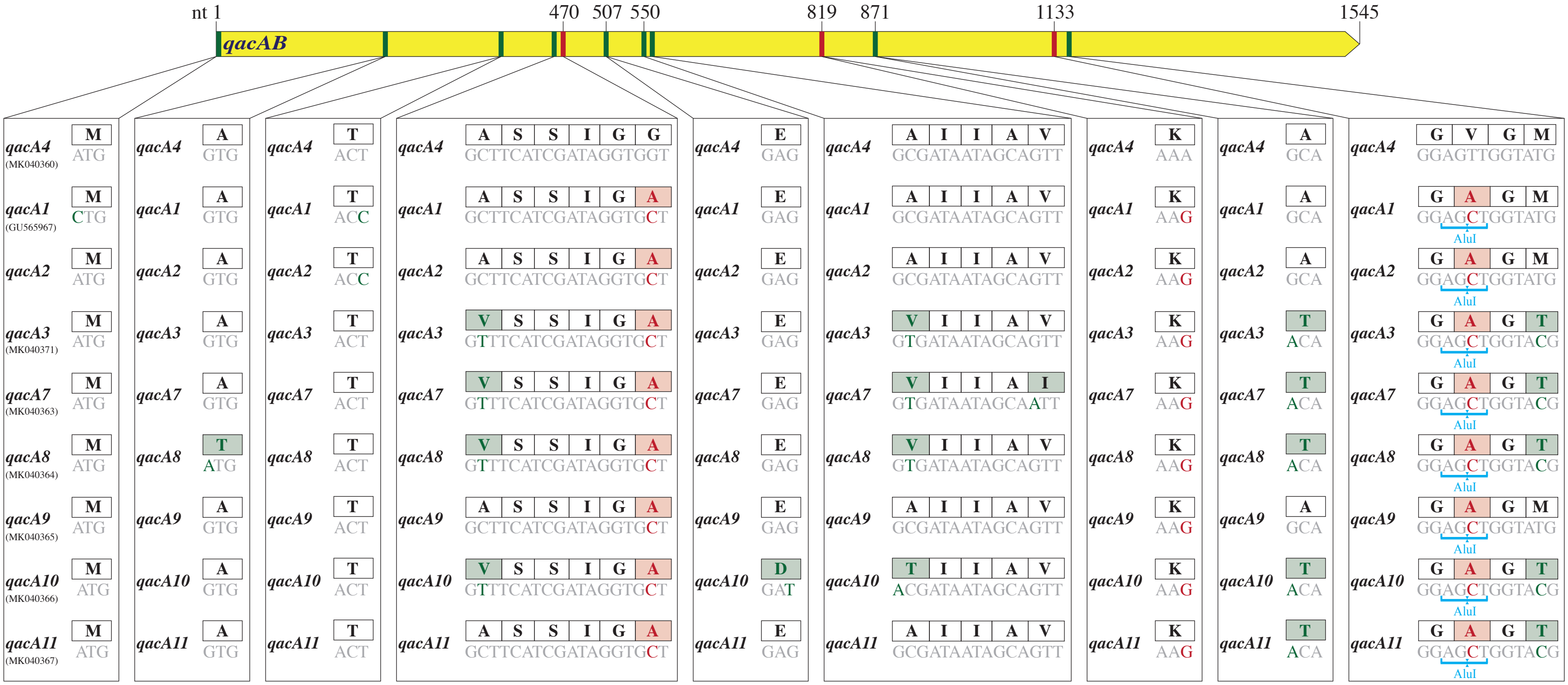


**Figure S1.** Comparison of the sequences of *qacA4* (MK040360), the 3 previously described *qacA* alleles (*qacA1*, *qacA2*, *qacA3*; Alam et al., 2003), and the 5 novel *qacA* alleles identified in the *qacA*-positive control isolates without elevated CHG MICs sequenced in this study. The associated AluI restriction sites are shown below the nucleotide sequences and the corresponding amino acid sequences are displayed in the boxes above the nucleotide sequences. The nucleotides which distinguish *qacA4* from all other alleles are highlighted in red. Those which distinguish one or more of these alleles from *qacA4* are displayed in green.

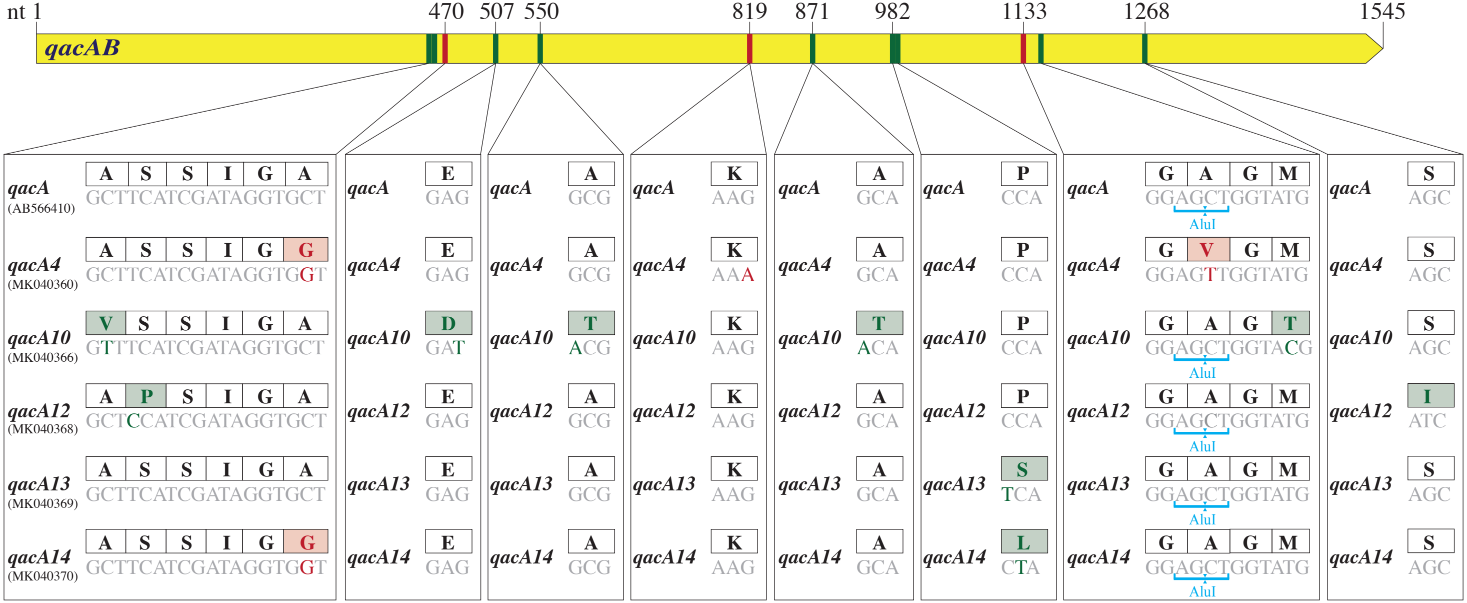


**Figure S2.** Comparison of the sequences of the reference *qacA* (AB566410), *qacA4* (MK040360), and the 4 novel *qacA* alleles identified in the *qacA*-positive isolates with elevated CHG MICs. The associated AluI restriction sites are shown below the nucleotide sequences and the corresponding amino acid sequences are displayed in the boxes above the nucleotide sequences. The nucleotides which distinguish *qacA4* from the reference *qacA* are highlighted in red. Those which distinguish the novel alleles from *qacA4* and the reference *qacA* are displayed in green.
