## Supplementary material for "A novel *qacA* allele results in an elevated chlorhexidine glu conate minimum inhibitory concentration in cutaneous *Staphylococcus epidermidis* isolates"

**Table S1.** BioSample accession numbers and phenotypic data for the isolates sequenced in this study. The CHG MIC is measured in μg/mL. “R” denotes the isolate is resistant to the specified antimicrobial and “S” indicates the isolate is susceptible to the specified antimicrobial.

| **Isolate ID** | **BioSample Accession No.** | ***qacA/B* RFLP** | ***qacA* allele** | ***smr*** | **CHG MIC** | **FOX** | **ERY** | **CIP** | **GEN** | **SXT** | **LZD** | **RIF** | **VAN** |
| --- | --- | --- | --- | --- | --- | --- | --- | --- | --- | --- | --- | --- | --- |
| *The initial 10 qacAB_273_-positive isolates with elevated CHG MICs randomly selected for whole-genome sequencing* | | | | | | | | | | | | | |
| 53.1 | SAMN07839754 | *qacAB_273_* | *qacA4** | No | 4 | R | R | R | R | R | S | S | S |
| 73.1 | SAMN07839755 | *qacAB_273_* | *qacA4* | Yes | 4 | R | R | R | R | R | S | S | S |
| 87.7 | SAMN07839756 | *qacAB_273_* | *qacA4* | Yes | 4 | R | R | R | R | R | S | S | S |
| 91.2 | SAMN07839757 | *qacAB_273_* | *qacA4* | Yes | 4 | R | R | R | S | R | S | S | S |
| 98.1 | SAMN07839758 | *qacAB_273_* | *qacA4* | Yes | 4 | R | R | R | R | R | S | S | S |
| 101.1 | SAMN07839759 | *qacAB_273_* | *qacA4* | Yes | 4 | R | R | R | R | R | S | S | S |
| 107.2 | SAMN07839760 | *qacAB_273_* | *qacA4* | No | 4 | R | R | R | R | R | S | S | S |
| 123.3 | SAMN07839761 | *qacAB_273_* | *qacA4* | Yes | 4 | R | R | R | R | R | S | S | S |
| 134.11 | SAMN07839762 | *qacAB_273_* | *qacA4* | Yes | 4 | R | R | R | R | R | S | S | S |
| 169.1 | SAMN07839763 | *qacAB_273_* | *qacA4* | Yes | 4 | R | R | R | R | R | S | S | S |
| *The 10 qacA-positive isolates without elevated CHG MICs randomly selected as positive controls* | | | | | | | | | | | | | |
| 20.1 | SAMN07839764 | *qacA* | *qacA3* | No | 1 | R | R | S | S | R | S | S | S |
| 66.3 | SAMN07839765 | *qacA* | *qacA7* | No | 2 | R | S | R | S | R | S | S | S |
| 68.5 | SAMN07839766 | *qacA* | *qacA8* | No | 2 | S | R | S | S | R | S | S | S |
| 97.1 | SAMN07839767 | *qacA* | *qacA8* | No | 2 | S | R | S | S | S | S | S | S |
| 99.1 | SAMN07839768 | *qacA* | *qacA9* | No | 1 | R | R | S | S | R | S | S | S |
| 106.1 | SAMN07839769 | *qacA* | *qacA9* | No | 2 | S | R | S | S | S | S | S | S |
| 110.3 | SAMN07839770 | *qacA* | *qacA10* | No | 2 | R | S | R | R | R | S | S | S |
| 128.1 | SAMN07839771 | *qacA* | *qacA10* | No | 1 | S | S | S | S | S | S | S | S |
| 131.1 | SAMN07839772 | *qacA* | *qacA9* | No | 2 | R | R | R | R | R | S | S | S |
| 135.2 | SAMN07839773 | *qacA* | *qacA11* | No | 1 | R | R | S | R | R | S | S | S |
| *The 4 qacB-positive isolates randomly selected as positive controls* | | | | | | | | | | | | | |
| 54.2 | SAMN07839774 | *qacB* |  | No | 1 | S | S | S | S | S | S | S | S |
| 81.4 | SAMN07839775 | *qacB* |  | No | 1 | R | S | S | S | S | S | S | S |
| 156.3 | SAMN07839776 | *qacB* |  | Yes | 2 | S | R | S | S | S | S | S | S |
| 174.1 | SAMN07839777 | *qacB* |  | No | 1 | R | R | S | S | S | S | S | S |
| *The 2 qacAB_273_-positive isolates without elevated CHG MICs* | | | | | | | | | | | | | |
| 36.5 | SAMN10232702 | *qacAB_273_* | *qacA5* | No | 1 | R | S | R | S | R | S | S | S |
| 125.10 | SAMN10232703 | *qacAB_273_* | *qacA6* | No | 0.5 | S | R | S | S | S | S | S | S |
| *The 9 qacA-positive isolates with elevated CHG MICs* | | | | | | | | | | | | | |
| 15.1 | SAMN10232693 | *qacA* | *qacA12* | No | 4 | R | R | R | R | R | S | S | S |
| 36.3 | SAMN10232698 | *qacA* | *qacA13* | No | 4 | R | R | R | R | R | S | S | S |
| 36.4 | SAMN10232699 | *qacA* | *qacA13* | No | 4 | S | R | R | R | R | S | S | S |
| 39.7 | SAMN10232694 | *qacA* | *qacA12* | No | 4 | R | R | R | R | R | S | S | S |
| 86.4 | SAMN10232700 | *qacA* | *qacA10* | No | 4 | R | R | S | S | S | S | S | S |
| 96.5 | SAMN10232701 | *qacA* | *qacA14* | Yes | 4 | R | R | R | R | R | S | S | S |
| 125.1 | SAMN10232695 | *qacA* | *qacA12* | No | 4 | R | R | R | R | R | S | S | S |
| 125.3 | SAMN10232696 | *qacA* | *qacA12* | No | 4 | R | R | I | R | R | S | S | S |
| 125.8 | SAMN10232697 | *qacA* | *qacA12* | No | 4 | R | R | R | R | R | S | S | S |
| *The isolate cured of qacA4* | | | | | | | | | | | | | |
| 107.2_cured_ | SAMN10490861 | Negative |  | No | 1 | R | R | R | R | R | S | S | S |
| *The 5 qacAB_273_-positive isolates with discordant susceptibility patterns (susceptible to cefoxitin, gentamicin, or erythromycin)* | | | | | | | | | | | | | |
| 73.5 | SAMN10232704 | *qacAB_273_* | *qacA4* | No | 4 | R | R | I | S | R | S | S | S |
| 134.15 | SAMN10232705 | *qacAB_273_* | *qacA4* | Yes | 4 | S | R | R | R | R | S | S | S |
| 171.1 | SAMN10232706 | *qacAB_273_* | *qacA4* | Yes | 4 | R | S | R | R | R | S | S | S |
| 171.10 | SAMN10232707 | *qacAB_273_* | *qacA4* | Yes | 4 | R | S | R | R | R | S | S | S |
| 173.6 | SAMN10232708 | *qacAB_273_* | *qacA4* | Yes | 4 | R | S | R | R | R | S | S | S |
| *A qacAB_273_-positive isolate collected from a previous chlorhexidine gluconate bathing study conducted at Seattle Children’s Hospital (Soma et al, 2012)* | | | | | | | | | | | | | |
| 13A1 | SAMN10237431 | *qacAB_273_* | *qacA4* | Yes | 4 | R | R | R | R | R | S | S | S |

* Isolate 53.1 contained two *qacA* alleles, with one being *qacA4*.

**Table S2.** GenBank accession numbers of the previously characterized *qacA* alleles and the 11 novel *qacA* alleles identified in this study.

| ***qacA* Allele** | **GenBank Accession No.** | **First reported** |
| --- | --- | --- |
| *qacA1* | GU565967 | (Alam et al., 2003). Previously described as “qacA prototype.” |
| *qacA2* | N/A | (Alam et al., 2003). Previously described as “qacA-V1.” |
| *qacA3* | MK040371 | (Alam et al., 2003). Previously described as “qacA-V2.” |
| *qacA4* | MK040360 | This study. |
| *qacA5* | MK040361 | This study. |
| *qacA6* | MK040362 | This study. |
| *qacA7* | MK040363 | This study. |
| *qacA8* | MK040364 | This study. |
| *qacA9* | MK040365 | This study. |
| *qacA10* | MK040366 | This study. |
| *qacA11* | MK040367 | This study. |
| *qacA12* | MK040368 | This study. |
| *qacA13* | MK040369 | This study. |
| *qacA14* | MK040370 | This study. |
